# A rapid alkalinization factor-like peptide EaF82 impairs tapetum degeneration during pollen development through induced ATP deficiency

**DOI:** 10.1101/2022.08.11.503650

**Authors:** Chiu-Yueh Hung, Farooqahmed S. Kittur, Keely N. Wharton, Makendra L. Umstead, D’Shawna B. Burwell, Martinique Thomas, Qi Qi, Jianhui Zhang, Carla E. Oldham, Kent O. Burkey, Jianjun Chen, Jiahua Xie

## Abstract

In plants, timely degeneration of tapetal cells is essential for providing nutrients and other substances to support pollen development. Rapid alkalinization factors (RALFs) are small, cysteine-rich peptides known to be involved in various aspects of plant development and growth, and defense against biotic and abiotic stresses. However, the functions of most of them remain unknown, while no RALF has been reported to involve in tapetum degeneration. In this study, we demonstrated that a novel cysteine-rich peptide EaF82 isolated from shy-flowering ‘Golden Pothos’ plants is a RALF-like peptide and displays alkalinizing activity. Its heterologous expression in Arabidopsis delayed tapetum degeneration and reduced pollen production and seed yields. RNAseq, RT-qPCR and biochemical analyses showed that overexpressing *EaF82* down-regulated a group of genes involved in pH changes, cell wall modifications, tapetum degeneration and pollen maturation as well as seven endogenous Arabidopsis *RALF* genes, and decreased proteasome activity and ATP levels. Yeast two-hybrid screening identified AKIN10, a subunit of energy-sensing SnRK1 kinase, to be its interacting partner. Our study reveals a possible regulatory role for RALF peptide in tapetum degeneration and suggests that EaF82 action may be mediated through AKIN10 leading to the alteration of transcriptome and energy metabolism, thereby causing ATP deficiency and impairing pollen development.

## Introduction

Since the discovery of systemin in 1991^1^, numerous plant peptide hormone-like sequences have been identified^2^ and the quest for understanding their functions and action modes is growing. One such peptide family is rapid alkalinization factors (RALFs), which are small cysteine-rich peptides (CRPs) widely present in plant kingdom^3^. Many RALF-like sequences have been identified from genomic sequences^4, 5^; and a phylogenetic analysis of 795 RALFs from 51 species showed that they fall into four distinct clades^5^. However, only a handful of RALFs belonging to clades I, II and III have been functionally characterized and shown to be involved in cell expansion, root and root hair development, pollen tube elongation/rupture and immunity, while the majority of them have not been studied functionally^6^. The clade IV is the least investigated and distinct from other clades with greater evolutionary divergence^5, 6^.

Studies related to the function and mechanism of RALFs are attracting greater attention due to their importance in plant growth and development, and defense against biotic and abiotic stresses^7, 8^. Recent studies on Arabidopsis AtRALF1/23/4/19 demonstrated that their actions are perceived through interactions with *Catharanthus roseus* Receptor Like Kinase 1 Like (CrRLK1L) receptors^6^. Binding of AtRALFs to CrRLK1L receptors triggers downstream signal transduction including down-regulation of H^+^-ATPase resulting in extracellular pH increase. The increased extracellular pH was thought to strengthen the cell wall to prevent pathogen invasion and make alkaline apoplasts an unfavorable environment for pathogens^9^. The pH increase also counteracts the acidification-induced loosening of cell wall leading to inhibition of root growth and pollen tube elongation^10^. As for clade IV RALFs, many members are thought to be involved in pollen development based on their transcriptional patterns. Studies on male sterile Chinese cabbage lines found that three out of 14 identified differentially expressed *RALFs* belong to clade IV^11^. Studies on four pollenless Arabidopsis tapetum mutants also revealed that five out of seven differentially expressed *RALFs* are members of clade IV^12^. In Arabidopsis, six out of eight pollen-abundant *AtRALFs* (4/8/9/15/19/25/26/30) were found to belong to clade IV^13^. However, to date, their precise roles and underlying mechanism of action in pollen development are still unknown.

We previously isolated a novel gene *EaF82* from shy-flowering ‘Golden Pothos’ plants (*Epipremnum aureum*)^14, 15^, encoding a 120-amino acid CRP^16^. ‘Golden Pothos’ is one of the most popularly grown houseplants worldwide with variegated leaves and can be used for studying chloroplast biogenesis by comparing genes in the cells between color defective and normal green sectors^14, 16^. *EaF82* was found to be expressed in all types of vegetative tissues, such as leaves, petioles, nodes and roots at different levels^14^. It was differentially expressed between green and yellow/white sectors of variegated ‘Golden Pothos’ leaves^14, 16^, and its expression was positively regulated by auxin^16^. However, EaF82 function remained unknown, and its expression in the reproductive organs has not been investigated because all *E. aureum* plants do not flower^15^.

In the present study, we show that EaF82 is closely related to clade IV RALFs and the synthetic EaF82 peptide can induce alkalinization of tobacco suspension cell culture medium as typical RALFs. To investigate its potential function as a clade IV RALF-like peptide involved in pollen development in the shy-flowering ‘Golden Pothos’ plants is challenging. Therefore, we adopted a gain-of-function approach commonly used for studying genes in non-model, long life cycle species (e.g. poplar)^17^ and ectopically expressed *EaF82* in model species Arabidopsis (*Arabidopsis thaliana*) and tobacco (*Nicotiana benthamiana*) which have no *EaF82* homologous gene^16^. Overexpressing *EaF82* in Arabidopsis plants was found to affect pollen production and seed-setting associated with induced transcriptomic changes and decreased ATP levels in flowers. Furthermore, AKIN10, a catalytic α-subunit of energy sensor - sucrose non-fermenting related kinase 1 (SnRK1), was identified as an interacting partner of EaF82. Its potential action mechanism through binding to AKIN10 to alter transcriptome and energy metabolism leading to impaired tapetum degeneration and pollen development is discussed.

## RESULTS

### EaF82 is a clade IV RALF-like peptide

Our previous search for EaF82 homologs using protein BLAST only found an antimicrobial peptide MiAMP1 from *Macadamia integrifolia* as the closest match with E-value of 0.094 and 48% similarity^16^. In the current study, we further analyzed its sequence features and used a recent comprehensive peptide classification system based on peptide structural features and biological functions^8^ to find its closely related peptide homologs. In the deduced 120-amino acid EaF82 peptide sequence from the cloned cDNA, a 30-amino acid signal peptide (Fig. 1a) was predicted by SignalP 5.0^18^. EaF82 contains four cysteines at positions 42, 54, 81 and 95 with potential to form two intramolecular disulfide bridges (Fig. 1a). Its primary structural features were found to be most similar to a group of CRPs classified as “nonfunctional precursor” by Tavormina et al.^8^. The number of cysteines and the amino acid patterns around disulfide bonds are known to be conserved for proteins having similar folding and function, and can be used as the basis for protein classification^19^. Thus, these criteria were used for further classification and revealed that the features of EaF82 were closest to those of RALFs among the listed CRPs, having two predicted intramolecular disulfide bridges and an *N*-terminal signal peptide^20^.

**Fig. 1.**
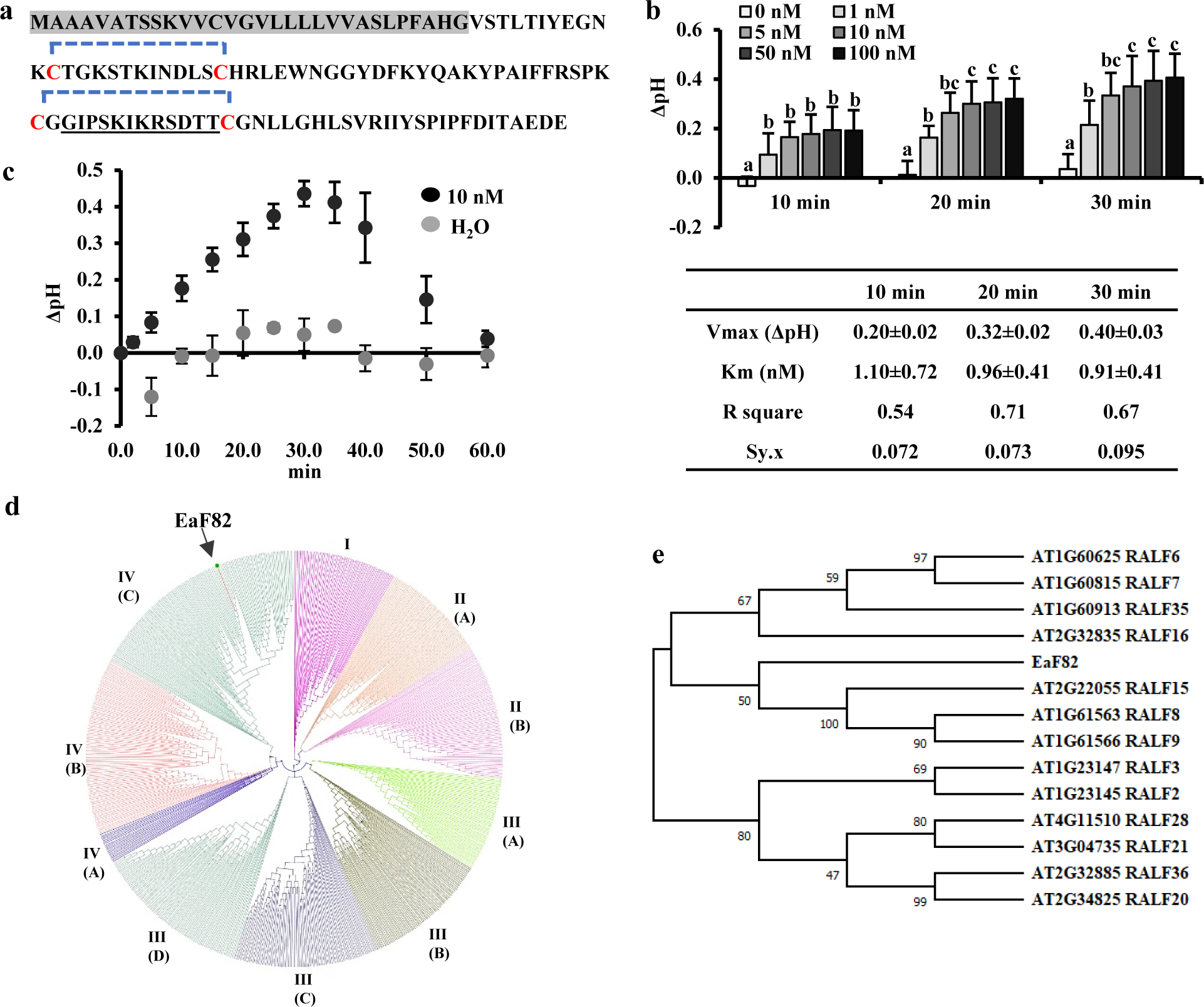
EaF82 peptide and alkalinization assay. **a** Amino acid sequence. The predicted signal peptide is highlighted in gray. Four cysteines (C) are marked in red with predicted potential intramolecular disulfide bridges indicated with blue brackets. Sequences for making antibody is underlined. **b** Alkalinization assays of EaF82-S (EaF82 without signal peptide). Six different concentrations (0, 1, 5, 10, 50, and 100 nM) were tested and the pH changes (Δ pH) were measured after 10, 20 and 30 min. Data plotted were the average of five independent experiments ± SD. The Km and Vmax are listed below. Data marked with the same letter are not significantly different by the LSD test at 5% level of significance. **c** Alkalinization activity measured for 60 min. Data represent the average of five independent experiments ± SD. **d** Phylogenetic tree of EaF82 among 795 RALFs from 51 plant species. Clade I, II, III and IV as well as their subgroups are categorized following Campbell and Turner^5^. **e** Phylogenetic tree of EaF82 with clade IV-C of AtRALFs.

The hallmark of RALFs is their ability to rapidly alkalinize tobacco cell culture media upon addition of exogenous peptide^7^. To investigate whether EaF82 also possesses a RALF-like alkalinizing activity, EaF82 peptide without signal peptide was synthesized (designated as EaF82-S) and its activity was measured, including kinetic parameters *Km* and Vmax. EaF82-S exhibited alkalinizing activity with a Vmax of delta pH ∼0.4, which was about half of reported values^7^, and *Km* of 1 nM (Fig. 1b) close to reported values for RALFs^7, 10^. The pH increase peaked at 30 min and returned to baseline after 60 min (Fig. 1c). In contrast, the EaF82 alkalinizing activity was abolished when the peptide was reduced and alkylated to break disulfide bonds (a negative control) (Supplementary Fig. 1a). The results indicate that EaF82 has a RALF-like alkalinizing activity.

To examine the evolutionary relationship of EaF82 with RALFs, phylogenetic analysis was performed using the method and sequence information described by Campbell & Turner ^5^. We found that EaF82 belonged to clade IV-C (Fig. 1d). Since Arabidopsis was to be used to heterologously express *EaF82* for functional studies, we further used 13 out of 14 Arabidopsis clade IV-C RALFs from previous publication^5^ to align with EaF82 by MUSCLE^21^ and analyze their evolutionary relationships using MEGA X^22^. The previously reported AtRALF17 (AT2G32890)^5^ was excluded as it lacks common features of RALF family members and is very likely not a RALF peptide^23^. Phylogenetic analysis revealed that EaF82 was closely related to AtRALF8/9/15 (Fig. 1e), which are known to be abundantly expressed in pollen^13^.

### EaF82 is expressed and accumulated in anthers but not in pollen

To investigate its physiological function, a reporter gene *sGFP*(S65T) in-frame with *EaF82* driven by *EaF82* promoter (*EaF82p::EaF82-sGFP* designated as TA) or by a constitutive *CaMV 35S* promoter (*35Sp::EaF82-sGFP* designated as TB) (Supplementary Fig. 2) was transformed into Arabidopsis. The former was used to determine the *EaF82* expression sites while both of them were used to investigate the effects of EaF82 peptide on Arabidopsis growth and development with a special attention to pollen development. When the *EaF82* transcriptional and translational expression levels in TA transgenic seedlings were analyzed by RT-PCR and immunoblotting, expected sizes of PCR and protein products were detected (Supplementary Fig. 3a). These validated TA transgenic seedlings showed GFP signals in roots, especially at the basal and apical meristems of primary and lateral roots (Supplementary Fig. 3b, c), where auxin is known to be accumulated^24^. The results are consistent with our previous finding in which GUS was found to accumulate at the same sites when driven by *EaF82* promoter^16^. GFP signals were observed in anthers and filaments of mature flowers (opened), but only occasionally in released pollen grains (Fig. 2a, b). We also checked the GUS activity in our previously created *EaF82p::GUS* plants^16^. Similarly, less GUS activity was observed in mature flowers than in early stages of closed flower buds (Fig. 2c), where high auxin accumulation has been reported^25^.

**Fig. 2.**
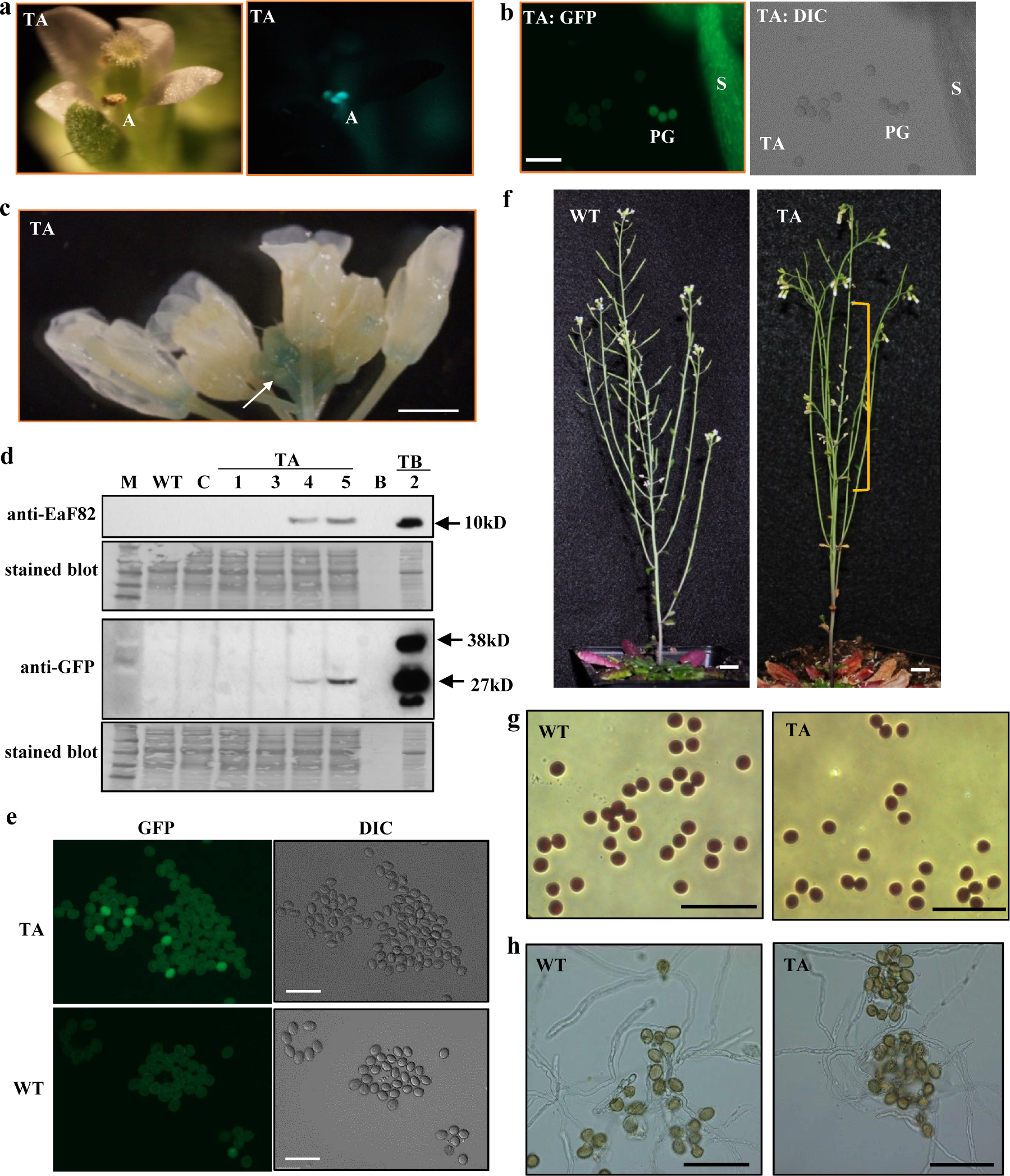
The functional characterization of Arabidopsis transgenic lines carrying *EaF82* promoter driving *EaF82-sGFP* (TA) or *GUS*. **a** The newly opened flower (left) shows GFP signal in anthers (right). A: anther. **b** Confocal microscopy shows GFP (left) and DIC (differential interference contrast, right) in stamen and some pollen grains. S: stamen; PG: pollen grains; Bar = 100 µm. **c** GUS staining of the flowers of *EaF82p::GUS*. White arrow indicates GUS activity at the early developmental stages of flowers. Bar = 1.5 mm. **d** Immunoblot of pollen proteins against anti-EaF82 and anti-GFP antibodies. Stained blots show protein loading. TA-1, -3, -4 and -5: four independent lines. TB-2: Proteins isolated from flowers of Arabidopsis transgenic *35Sp::EaF82-sGFP*. B: blank. C: vector control. WT: wild-type. M: protein size marker. **e** Confocal microscopy detecting GFP in collected pollen grains from Arabidopsis transgenic *EaF82p::EaF82- sGFP* (TA) plants. WT is a negative control. Bar = 100 µm. **f** A 10-week old TA plant (right) bears aborted siliques in primary inflorescence stalk (yellow bracket) compared to normal WT (left). Bar = 1 cm. **g** Collected pollen grains from fully opened flowers stained with iodine-potassium iodide. **h** Germinated pollen under germination medium. Bar = 100 µm.

EaF82 is closely related to AtRALF8/9/15 (Fig. 1e), which are reported to be abundant in pollen^13^. However, the expression of *sGFP* driven by *EaF82* promoter was rarely detected in pollen grains (Fig. 2b). Thus, its protein level in pollen was further examined in detail. Seedlings with confirmed EaF82 accumulation from four independent TA lines (Supplementary Fig. 4a) were transferred to soil and grew into mature plants to collect pollen grains. These TA seedlings grew normally on MS medium (Supplementary Fig. 4b) and became mature plants on soil, implying that EaF82 might have no effect on vegetative growth as reported for AtRALF1/8/23^6^. Collected pollen grains had little detectable EaF82 and GFP proteins as revealed by immunoblotting (Fig. 2d). Consistent with this observation, under the confocal microscope only a few pollen grains displayed GFP signals (Fig. 2e). These results indicate that EaF82 was not abundant in pollen. Likely, the observed GFP signals in anthers (Fig. 2a) could come from the surrounding tissues instead of pollen grains.

### Overexpressing EaF82 affected pollen development and seed-setting

Interestingly, the above TA lines showed many unpollinated pistils and short siliques on the primary inflorescence stalks (Fig. 2f). This observation raised a question of whether EaF82 plays a role in seed-setting since a group of RALFs has been shown to affect the rate of seed-setting through inhibiting pollen tube elongation^10^. We collected pollen even though much less pollen grains produced in some flowers, and then examined pollen viability and pollen tube elongation. There were no differences in the general features of harvested pollen grains (Fig. 2g) as well as no obvious differences in pollen germination ability, nor *in vitro* pollen tube elongation (Fig. 2h) compared to those of wild-type (WT). Therefore, the effects of EaF82 on pollen viability are likely minimum once pollen is produced. These results led us to speculate that those unpollinated pistils and short siliques on the primary inflorescence stalks could be the results of high EaF82 accumulation in the surrounding tissues of pollen sacs causing either no pollen or less pollen available for the pollination at the specific stage of development. Since the TA lines were driven by an auxin responsible *EaF82* promoter^16^, it is possible that the accumulation of EaF82 along the inflorescence stalks was uneven due to the uneven auxin distribution affected by the growth conditions.

The above speculation that high EaF82 accumulation in the surrounding tissues of pollen sacs might affect pollen development and production was supported by the observations from TB lines with the *EaF82* driven by a strong constitutive *CaMV 35S* promoter. First, although the same floral dipping procedure for creating TA lines was used, our transformation efforts with the TB cassette resulted in only two independent lines (TB-1 and TB-2), despite several attempts. The difficulty of TB transformation seemingly implies that the presence of high EaF82 might cause the developmental problem in transformed pollen and subsequently affect the production of transformed seeds. When two TB lines were grown on MS medium, their seedlings were normal as the WT (Supplementary Fig. 5a). Unfortunately, EaF82 and GFP were detected only in TB-2 line by immunoblotting (Supplementary Fig. 5b). TB-2 plants bore extremely few pollen grains and produced only 1-2 small siliques per plant even though they produced many flowers (Fig. 3a, b). TB-2 line exhibited more severe seed abortions compared to TA lines (Fig. 2f). In order to obtain ∼100 siliques, 72 independent plants were planted per subline of TB-2a and TB-2b. The average numbers of seeds per silique in TB-2 plants were reduced to only 4 compared to 49 in the case of WT (Fig. 3c).

**Fig. 3.**
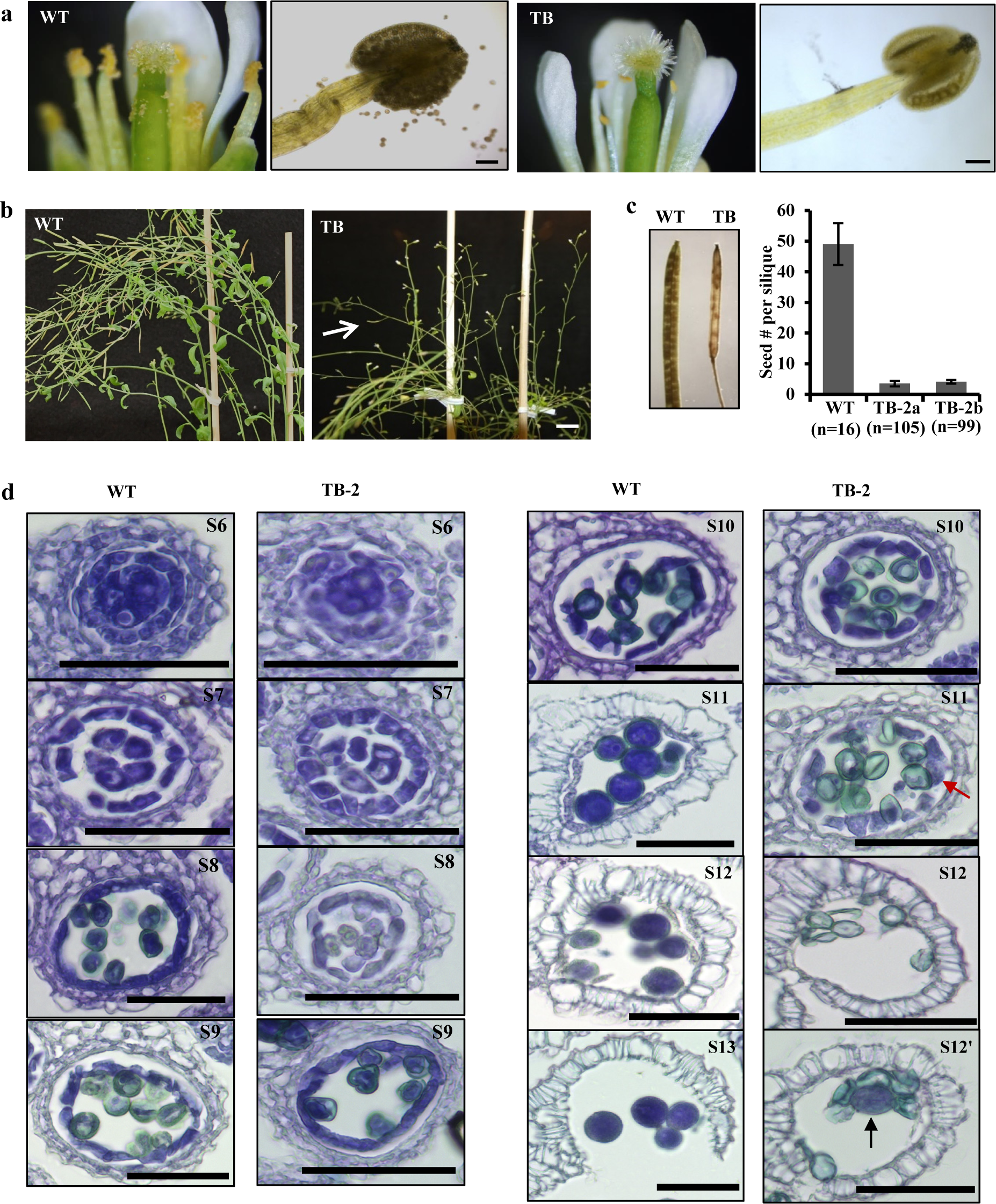
Male gametophyte and pollen development of Arabidopsis transgenic *35Sp::EaF82-sGFP* lines (TB) comparing to wild-type (WT). **a** Fully opened flower (left) and stained anther with iodine- potassium iodide (right) of WT and TB line, respectively. Bar = 100 µm. **b** TB plants (right) produce no siliques with occasionally observed small silique (white arrow) compared to WT (left). Bar = 1 cm. **c** TB silique carries less seeds than that of WT (left). The numbers of seeds per silique are plotted as the mean ± SD (right). n: numbers of siliques. **d** Male gametophyte development of TB line compared to WT. At the stage S11, the undegenerated tapetum (red arrow) was observed in TB line. At the stages S11 and S12, underdeveloped pollen was stained in light green, while mature pollen is in dark blue that occasionally was observed in TB line (black arrow). Bar = 50 µm.

Since the major difference between TA and TB lines was the promoters, we further tested promoter activities using multiple independent tobacco (*Nicotiana tabacum*) transgenic lines carrying genetic cassettes *35Sp::GUS* or *EaF82p::GUS* (Supplementary Fig. 2) to compare their *GUS* transcript levels. RT-qPCR results showed ∼3.5-fold higher *GUS* expressions in transgenic *35Sp::GUS* plants than those of transgenic *EaF82p::GUS* (Supplementary Fig. 6), indicating that the activity of *CaMV 35S* promoter is stronger than that of *EaF82* promoter. Thus, the severe seed abortion phenotype in TB-2 driven by the *CaMV 35S* promoter probably could result from constitutively higher expression of EaF82-sGFP compared to TA lines driven by the *EaF82* promoter, supporting that the high expression levels of EaF82 may be responsible for the pollenless and reduction of seed yield.

### Overexpressing EaF82 delayed tapetum degeneration during pollen development

The observed pollenless phenotype in TB-2 (Fig. 3a) suggested that the pollen development was compromised. Considering the observed accumulation of EaF82 in anthers as well as closed flower buds (Fig. 2a-c) where pollen is under development, we first investigated at which stage pollen development was impaired by histological analysis. In Arabidopsis, flower buds are clustered and the flower development can be divided into 12 developmental stages using a series of landmark events as described by Smyth et al.^26^. The anther development can be further divided into 14 stages based on the visualization of distinctive cellular events under microscope^27^. According to the key events of each stage, the pollenless TB-2 line was examined and found that stages of microspore mother cells undergoing meiosis and generating tetrads of haploid microspores, and microspores releasing from the tetrads (up to stage S8) were similar to those in WT (Fig. 3d). The abnormality was noticed at stages S11 and S12. At S11, the complete disappearance of tapetum occurred in WT but not in TB-2, and hence most of TB-2 pollen grains were aborted at S12 (Fig. 3d). These observations were further confirmed in another TB-2 flower cluster (Supplementary Fig. 7), indicating that EaF82 impairs tapetum degeneration and pollen development.

### EaF82 also affected pollen development and seed-setting in tobacco plants

To verify that the observed effects of EaF82 on pollen development and seed-setting were not limited to Arabidopsis, a genetic cassette *35Sp::EaF82* (designated as TC) (Supplementary Fig. 2) was expressed in *N. benthamiana* plants. Because tobacco leaf disc transformation method is efficient, this approach was used to generate 15 independent transgenic lines (T1-T15) with seven of them exhibiting a 3 to 1 segregation. Out of these seven lines, four (T2, T3, T8 and T11) had detectable EaF82 in leaves (Fig. 4a) and flowers (Fig. 4c) as revealed by immunoblotting. Mature anthers of these four transgenic lines were often shriveled in appearance with no or less pollen grains while those of the vector control lines were dehisced with many released pollen grains (Fig. 4b). Transgenic flowers were either not fertilized to develop seed pods or partially fertilized to develop small seed pods compared to those normal pods produced from the control lines (Fig. 4d). Among produced pods, the seed numbers per pod of lines T2, T3, T8 and T11 were reduced by 42%, 22%, 50% and 58%, respectively, compared to the control lines (Fig. 4e). The results from these transgenic lines indicate that EaF82 also inhibits pollen development and seed-setting in tobacco plants.

**Fig. 4.**
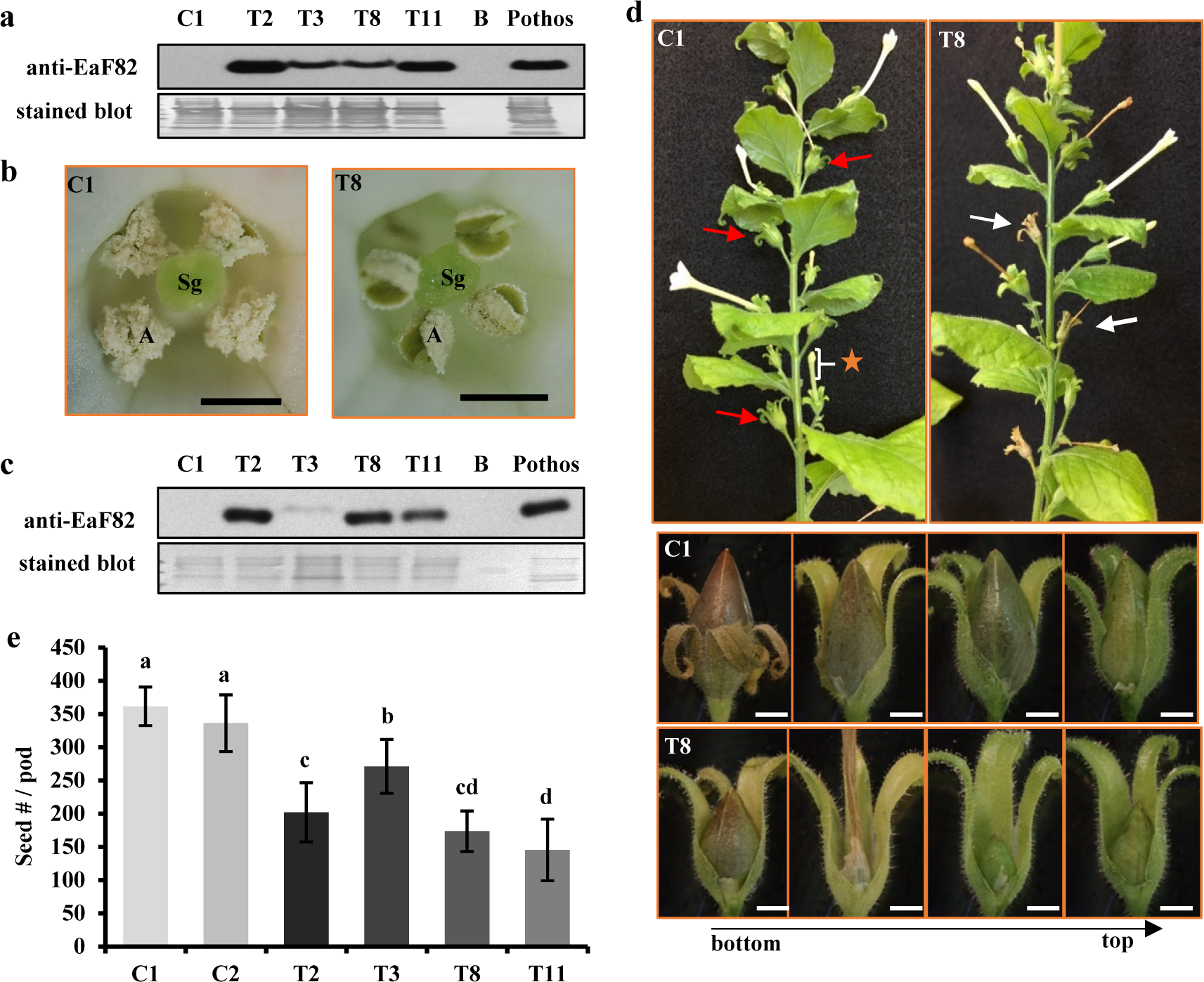
The functional characterization of tobacco transgenic *35Sp::EaF82* lines (T2, 3, 8 and 11) comparing to two independent vector control lines (C1 and C2). **a** Immunoblot of leaf proteins against anti-EaF82 antibody. Stained blot shows protein loading. **b** T8 (right) and C1 (left) opened flowers. Sg: stigma. Bar = 200 µm. **c** Immunoblot of unopened flower proteins against anti-EaF82 antibody. The unopened flowers are as indicated in (d) with orange star. **d** Transgenic plants with normal (red arrow) and aborted (white arrow) seed pods. Enlarged seed pods are shown below. **e** Seed counts per pod. Data plotted are the average from six independent plants per line using 10 seed pods per independent plant ± SD. Data marked with the same letter are not significantly different by the LSD test at 5% level of significance. Bar = 200 µm.

### Overexpressing EaF82 induced transcriptome changes

To elucidate the observed delayed tapetum degeneration during pollen development, RNAseq analysis was performed to gain molecular insights of EaF82 associated transcriptional alteration. To ensure the pollen abortion is associated with overexpressed EaF82 but not sGFP, Arabidopsis transgenic lines with the TC genetic cassette (*35Sp::EaF82*) without *sGFP* (Supplementary Fig. 2) were created. Similar to TB in which the high expression of EaF82 resulted in nearly no transformed seeds, transformation with TC genetic cassette was also challenging and produced only two lines (TC-1 and TC-2) with detectable EaF82 levels (Fig. 5a, b). They grew normally during the vegetative stage compared to the WT (Fig. 5c) but exhibited seed abortion with most undeveloped siliques (Fig. 5d) like the TB line. Their unopened flowers had detectable EaF82 (Fig. 5e) and later could develop a few viable pollen grains. When a few pollen grains were harvested for germinating, the pollen tube growth was normal (Fig. 5f, g). Among developed TC siliques, however, many of them were short in length compared to most WT siliques (Fig. 5h). A large-scale measurement using siliques from 10 primary stems per line showed that approximately 20.5% of TC-1 and 27.8% of TC-2 siliques did not develop longer than 0.3 cm (Fig. 5h), which contained no seed at all (Supplementary Fig. 8). Only 16.6% of TC-1 and 18.8% of TC-2 siliques were longer than 1.0 cm with some seeds while that of WT was 88.5% (Fig. 5h).

**Fig. 5.**
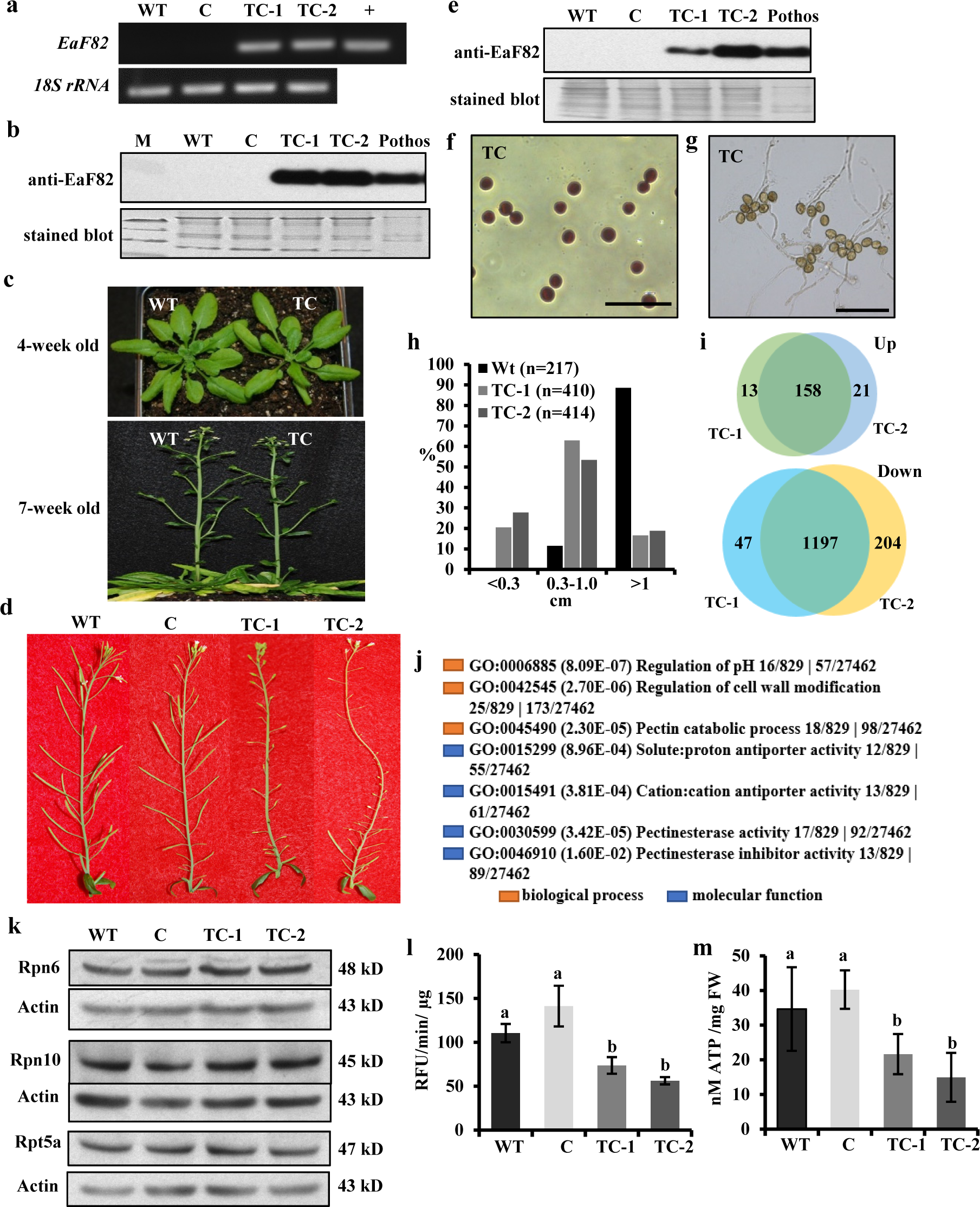
The functional characterization and RNAseq analysis of Arabidopsis transgenic *35Sp::EaF82* lines (TC-1 and -2) compared to wild-type (WT) and vector control (C). **a** RT-PCR of leaf tissues using primer pair specific to *EaF82* and *18S rRNA*. +: plasmid DNA. **b** Immunoblot of leaf proteins against anti-EaF82. Stained blot shows protein loading. M: protein size marker. **c** Normal growth of TC and WT plants. **d** Aborted siliques in TC-1 and TC-2. **e** Immunoblot of proteins from unopened whole flowers against anti-EaF82. **f** Stained pollen and **g** germinated pollen. The descriptions are the same as in Fig. 2. Bar=100 µm. **h** Histogram of the silique lengths. Data plotted are the percentages of total siliques. n: numbers of siliques. **i** The number of up- and down-regulated DEGs (≥ 2-fold) in TC-1 and -2 lines. **j** Enriched GO terms from down-regulated DEGs related to pH regulation and cell wall modification. **k** Immunoblots of three subunits of the proteasome in the early developmental flowers against anti-Rpn6, anti-Rpn10 and anti-Rpt5a antibodies showed no significant difference among all samples. **l** Proteasome activity. Data plotted are the average of three biological replicates ± SD. RFU: relative fluorescence units. **m** ATP content. Data plotted are the average of four biological replicates ± SD. FW: fresh weight.

To gain molecular insights into transcriptional changes during pollen development, the gene expression profiles of the unopened flower bud clusters covering all anther developmental stages from TC-1 and TC-2 lines were analyzed by RNAseq along with a vector control line. Two TC lines were used to perform double verification of any observed differentially expressed genes (DEGs). The numbers of reads were between 18.3 to 20.0 million (Supplementary Table 1), and listed genes reported as FPKM (Fragments Per Kilobase Million) were 28,296 (Supplementary Data Set 1), covering ∼90% of total nuclear, mitochondria and chloroplast genes (Supplementary Fig. 9a). The numbers of common DEGs in TC-1 and TC-2 either increased or decreased 2-fold with adjusted *p*-value for false discovery rate (FDR) < 0.05 compared to the vector control were 158 and 1197, respectively (Fig. 5i; Supplementary Table 2, 3). Hierarchical cluster analysis of 74 down-regulated DEGs with log_2_FC≤-1.5 (reduced ≥ 2.8-fold) plotted in a heatmap with R function showed a strong correlation between TC-1 and TC-2 lines (Supplementary Fig. 9b). The above results demonstrated that overexpressing *EaF82* induced ∼5% nuclear gene expression changes ≥ 2-fold in the early development of flowers.

### Overexpressing EaF82 suppressed the expression of genes involved in cell wall modifications and pH changes

To determine the affected pathways and related genes in overexpressing *EaF82* lines, identified 158 up- and 1197 down-regulated DEGs were subjected to Gene Ontology (GO) analysis using PANTHER v16^28^, and some selected DEGs were subjected to RT-qPCR validation. In the up-regulated DEGs, only regulation of developmental process (GO:0050793) in biological process and sequence-specific DNA binding (GO:0043565) in molecular function were significantly enriched with a similar group of genes (Supplementary Data Set 2), including flowering-related genes *AGL19, TFL1, AGL20 (SOC1), AGL24, AGL42, FD* and *SAP* (Supplementary Table 2)^29, 30^. All four up-regulated AGAMOUS-LIKE DEGs *AGL19, AGL20, AGL24* and *AGL42* as well as 27 down-regulated DEGs involved in four different categories (Table 1, 2) were selected for validation by RT-qPCR. Their transcriptional changes from RNAseq analysis were well confirmed by RT-qPCR results with similar fold decreases (Table 1, 2).

**Table 1.**
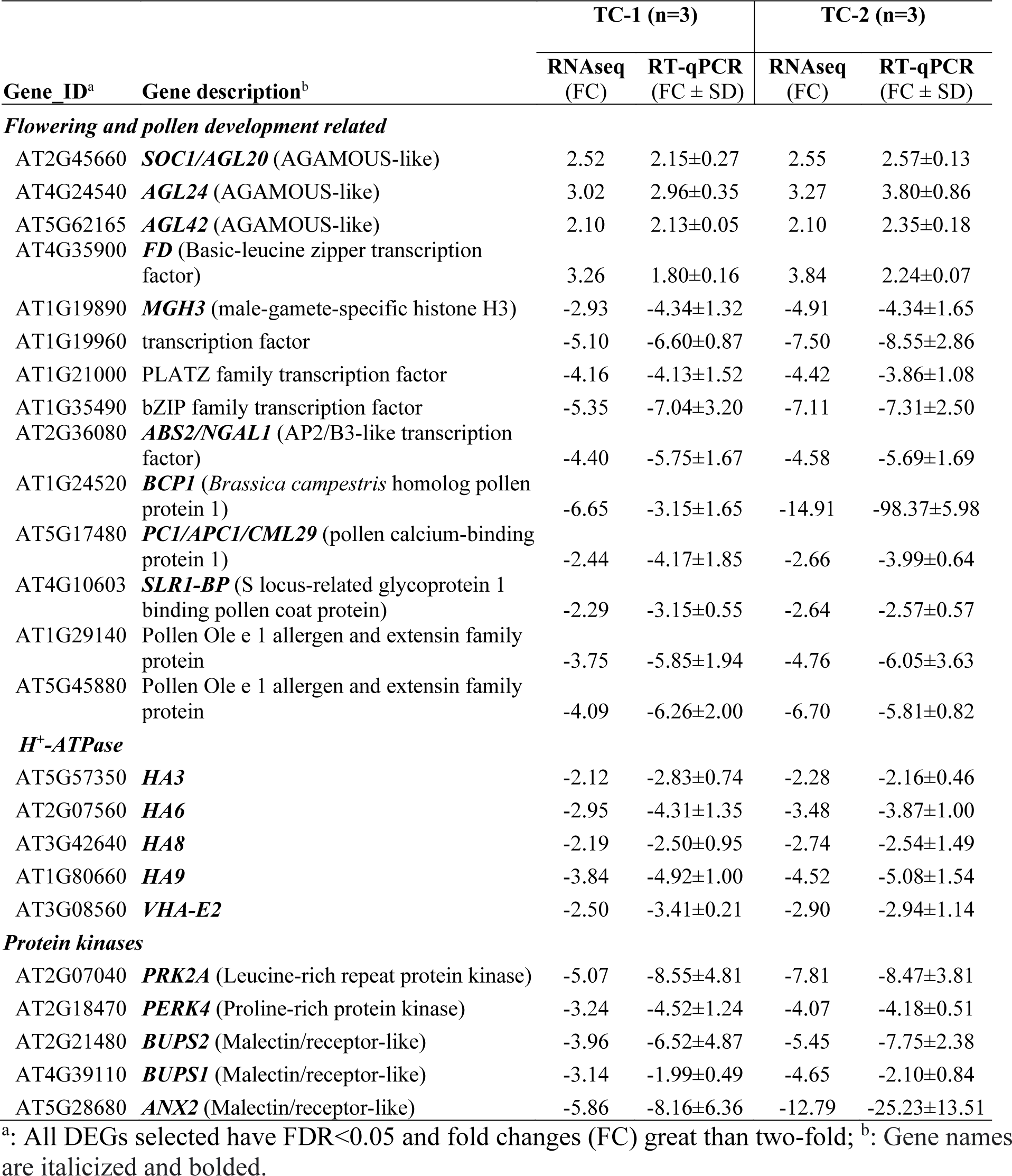
RT-qPCR of a subset of selected DEGs

**Table 2.** RT-qPCR of seven RALF genes and their predicted pI and molecular weight (MW)

| Gene_ID<sup>a</sup> | Gene description (clade)<sup>b</sup> | TC-1 (n=3) |  | TC-2 (n=3) |  | Predicted pI and MW<sup>c</sup> |  |
| --- | --- | --- | --- | --- | --- | --- | --- |
| | | RNAseq (FC) | RT-qPCR (FC ± SD) | RNAseq (FC) | RT-qPCR (FC ± SD) | pI | MW (kD) |
| AT1G28270 | ***RALF4*** (III-B) | -4.27 | -7.00±2.79 | -5.96 | -8.37±4.59 | 9.76 | 8.7 |
| AT1G61563 | ***RALF8*** (IV-C) | -3.67 | -5.26±1.60 | -5.35 | -5.66±1.00 | 9.30 | 6.6 |
| AT1G61566 | ***RALF9*** (IV-C) | -3.51 | -4.77±1.26 | -5.27 | -5.74±1.38 | 9.27 | 6.6 |
| AT2G22055 | ***RALF15*** (IV-C) | -5.73 | -2.48±0.39 | -23.41 | -16.92±2.40 | 9.76 | 6.5 |
| AT2G33775 | ***RALF19*** (III-B) | -5.26 | -9.10±5.00 | -9.69 | -8.14±3.43 | 10.57 | 7.5 |
| AT3G25165 | ***RALF25*** (IV-B) | -8.24 | -2.48±0.67 | -23.55 | -82.52±18.38 | 10.08 | 7.0 |
| AT4G14020 | RALF family protein (IV-A) | -3.39 | -4.29±1.30 | -3.78 | -4.02±1.27 | 10.17 | 6.8 |

<sup>a</sup>: All DEGs selected have FDR<0.05 and fold changes (FC) great than two-fold; <sup>b</sup>: Gene names are italicized and bolded; <sup>c</sup>: Amino acid sequences were retrieved from TAIR (https://www.arabidopsis.org/).

In the down-regulated DEGs, notably enriched GO terms related to the observed characteristics of transgenic plants were those involved in the regulation of pH, and cell wall modification (Fig. 5j; Supplementary Data Set 3). The overrepresented DEGs include those encoding 29 pectin methylesterases (PMEs)/pectin methylesterase inhibitors (PMEIs), along with five H^+^-ATPases and 16 cation/H^+^ antiporters involved in regulating pH changes (Table 1; Supplementary Table 4). These DEGs are known to play important roles in modulating the cell walls as an adaptation to stresses during plant development^31^. PME activity is regulated by pH changes through coordinating with H^+^-ATPases and cation/H^+^ antiporter^32^. Both PMEs and PMEIs were found to be highly expressed in flower buds and anthers^33, 34^. Among the down-regulated pollen specific *PMEs* (Supplementary Table 4), *PPME1* and *PME48* are regulated by RGA, a GA repressor DELLA^35^; while *PME5/VGD1*, *PME4/VGDH1* and *VGDH2* are regulated by MYB80, a transcription factor regulating both tapetum and pollen development^36^. These down-regulated DEGs together with reported roles of RGA and MYB80 at the late stage of tapetum degeneration and pollen development^35, 36^ suggest that cell wall modification was altered during the pollen development, supporting observed impaired tapetum degeneration and pollen abortion (Fig. 3d).

Moreover, seven DEGs (*AGP*5/6/11/14/23/24/40) encoding highly glycosylated arabinogalactan proteins (AGPs), which play key roles in pollen wall formation^37^, were also down-regulated (Supplementary Table 4). In Arabidopsis, pollen grains from *AGP6* and *AGP11* double mutants have been reported to exhibit pollen wall collapse^38^, and *AGP6*, *AGP11*, *AGP23* and *AGP40* are known to be involved in nexine formation^37^. These down-regulated *AGP*s present a strong correlation with the observed aborted pollen at stages S11 and S12 (Fig. 3d; Supplementary Fig. 7).

### EaF82 suppressed the expression of genes involved in tapetum degeneration and a group of *AtRALF*s

Pollen is developed inside the anther locules^39^. The innermost layer of anther locules is the tapetum, which is the main tissue providing nutrition and enzymes for pollen development and pollen wall formation^40, 41^. Mounting evidence suggests that the pollen developmental process is tightly linked to the development of tapetum^41, 42^ with the latter divided into three developmental stages: tapetum differentiation, tapetum formation, and tapetum degeneration through program cell death (PCD)^40^. Tapetal cells appear at the stage S5 and their degradation is initiated at the stage S10^27^. Our histological results of pollen development (Fig. 3d; Supplementary Fig. 7) showed that overexpressing EaF82 interrupted tapetum degeneration, but not tapetum differentiation and formation. This was supported by RNAseq data, which showed that many known genes involved in early stages of tapetum differentiation and tapetum formation^41^ were detected but not differentially expressed (Supplementary Table 5), whereas expression of *CEP1* involved in late stage of tapetum degeneration was found to be reduced ∼16-fold (Supplementary Tables 4, 5). *CEP1* encodes a papain-like cysteine protease and participates in tapetal cell wall hydrolysis^43^. Previous studies have shown that overexpression *CEP1* advanced the tapetal cell wall degeneration to early S7, while *cep1* mutant lacking functional CEP1 failed tapetum degeneration^43^. Generally, tapetum degeneration accompanies with tapetum cell wall degradation^43^. Therefore, ∼16-fold down-regulated *CEP1* together with a large number of cell wall modification associated genes (Supplementary Table 4) supports the observed impairment of tapetal cell degradation (Fig. 3d; Supplementary Fig. 7).

In addition to the above genes, seven *AtRALFs,* including *AtRALF*4/8/9/19/25 observed to be down-regulated in four tapetum mutants^12^, were also found in our down DEGs (Table 2; Supplementary Table 4). Their down-regulation was further confirmed by RT-qPCR showing similar fold reductions in their expressions (Table 2). These *AtRALF*s might be responsible for cell-to-cell communication between tapetal cells and pollen cells, an important process for pollen development. Their real functions however, have not been investigated yet^12^.

### Overexpressing EaF82 decreased proteasome activity and ATP levels

The tapetum degeneration is a process resembling apoptosis-like PCD^42^, and is essential for pollen development^44^. Pollen abortion has been observed with delayed tapetum PCD^45^. Both proteases and proteasomes are critical to the progression of tapetum PCD^44, 46^. Besides *CEP1*, there are seven additional downregulated DEGs encoding proteases (Supplementary Table 4), suggesting that PCD processes may be defective. Since no proteasome genes were found in our DEGs, we then examined the translational abundance of three subunits (Rpn6, Rpn10 and Rpt5) of the proteasome by immunoblotting and the proteasome activity in the early development of flowers. Among them, Rpt5a is one of the six AAA-ATPases of proteasome and essential for pollen development^47^. Immunoblotting showed no differences in their protein abundances (Fig. 5k). However, the proteasome activity in TC-1 and TC-2 lines was reduced by 33% and 49% compared to the WT, or by 48% and 60% compared to the vector control, respectively (Fig. 5l).

The results of reduced proteasome activity prompted us to further examine ATP levels because PCD is an energy dependent process and its initiation and execution are affected by ATP, Ca^2+^ and NO^48, 49^. Moreover, the proteasome assembly and activities are also ATP-dependent^50^. With the same tissues used for examining proteasome subunits and proteasome activity, the ATP levels were found to be indeed reduced by 38% and 57% compared to the WT, or 46% and 63% compared to the vector control, respectively (Fig. 5m). Their similar trends of reduction suggested that the reduced proteasome activity was likely the result of low ATP supply at the early developmental stages of flowers even though no DEGs directly linked to ATP production (such as genes coding for ATP synthases) were found. ATP serves not only as an intracellular energy molecule, but also as a signal molecule in the extracellular matrix of plant cells through coordinating with Ca^2+^and ROS^51, 52^. ATP defective mutants are male sterile^53, 54^, while elevated ATP increases seed yields^55^. ATP deficiency in TC lines is also consistent with many of observed down-regulated DEGs, which encode proteins directly or indirectly affected by ATP, such as six ATP-binding cassette (ABC) transporters and five H^+^-ATPases, as well as a group of ATP-binding receptor-like protein kinases (Table 1; Supplementary Table 4). These results suggest that overexpressing EaF82 lowered ATP levels, and triggered down-regulation of genes encoding for ATP-binding proteins which in turn affected PCD activity and resulted in delaying tapetum degeneration. Additionally, tapetum degeneration is essential for releasing nutrients to support pollen grain development and maturation^40, 41^. In the DEG, a large number of genes encoding transporters and transmembrane proteins for shuttling sugars, amino acids, and ions were also found to be down-regulated (Supplementary Table 4), inferring that the nutrient transport probably was limited to the developing pollen grains.

### AKIN10 is an interacting partner of EaF82

To understand how overexpressing EaF82 causes a decrease in ATP levels and down-regulation of so many genes, we conducted Y2H screening followed by 1-by-1 interaction validation assay with *EaF82-S* as the “bait” and a cDNA library made from Arabidopsis mitotic flower buds as the “prey” to identify EaF82 interacting partners. Using the PBS (Predicted Biological Score) system, a total of 46 EaF82 interacting proteins with score A, B, C or D were obtained as good candidates (Supplementary Table 6). Based on their biological functions and cellular components from published studies, seven candidates ABCF4, ALATS, FKBP-like peptidyl-prolyl cis-trans isomerase family protein, PAPP2C, TCH4, AKIN10 and SYTA (Supplementary Table 7) were selected to perform 1-by-1 Y2H validation assay. The assay results were shown in Supplementary Fig. 10 and summarized in Supplementary Table 8. Among the seven candidates, we found three: ABCF4, PAPP2C and AKIN10, to have the strongest interactions. ABCF4 (AT3G54540, also named as AtGCN4) is an ATPase and regulates plasma membrane H^+^-ATPase activity^56^, PAPP2C (AT1G22280) is a protein phosphatase^57^ while AKIN10 (AT3G01090) is a major cellular energy sensor in plants and orthologous to mammalian AMP-activated protein kinase (AMPK)^58^.

Given the key role of AKIN10 as the energy sensor in plants and its orthologous to mammalian AMPK^58^, the interaction between AKIN10 and EaF82 (Fig. 6a) could be critical and is likely to be responsible for observed ATP deficiency, and aberrant tapetum degradation. ABCF4 could be also involved in the EaF82-induced intracellular pH increase leading to cell wall modification. To elucidate the observed induced ATP deficiency (Fig. 5l), therefore, we focused on AKIN10 and performed co-IP analysis to validate its interaction with EaF82.

**Fig. 6.**
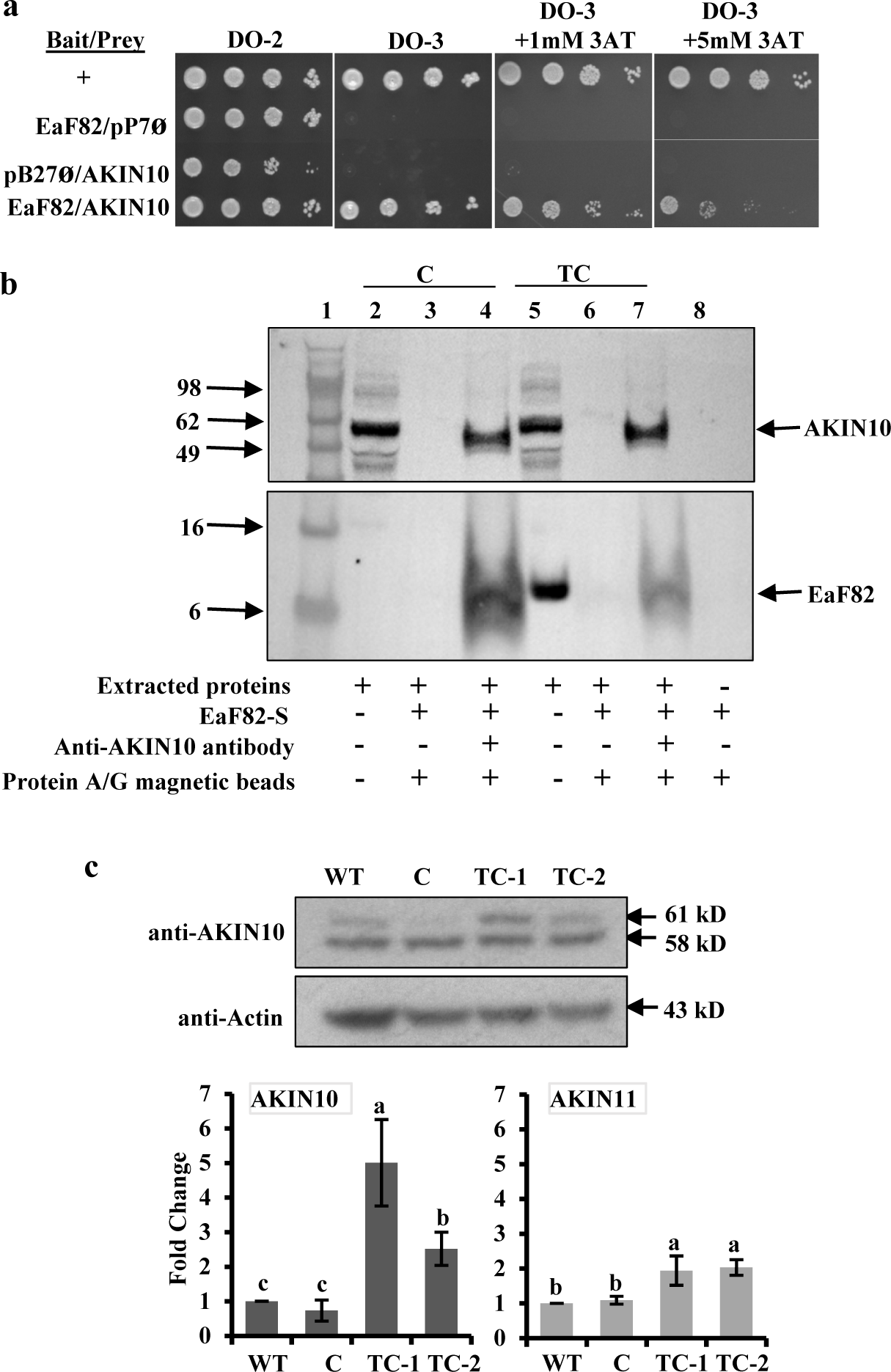
Identification and characterization of an EaF82 interacting partner. **a** Yeast growth tests of 1-by-1 Y2H assay on selective medium without (DO-2) or with (DO-3) histidine and 3- aminotriazole (3AT). Supporting information is detailed in Supplementary Fig. 10. +: positive interaction; pB27ø and pP7ø: empty vectors. **b** Co-immunoprecipitation (co-IP) analysis to validate the Y2H results of EaF82 and AKIN10 interaction. Protein extracts from floral tissues of vector control (C) and TC transgenic lines were spiked with (+) or without (-) EaF82-S. The complex was pull-downed with (+) or without (-) anti-AKIN10 antibody. The immobilized anti-AKIN10 on protein A/G magnetic beads could pull down EaF82 and AKIN10 complex from both C and TC lines (lanes 4 and 7). EaF82 was undetectable without immobilized anti-AKIN10 antibody (lanes 3 and 6). Lane 1: protein size marker. Lanes 2 and 5: extracted proteins from C and TC lines. Lane 8: EaF82-S alone bound to protein A/G magnetic beads. **c** A representative immunoblot of proteins from unopended flowers against anti-AKIN10 that detects both AKIN10 (61 kD) and AKIN11 (58 kD). The average of band intensity is plotted as three independent experiments ± SD (below). Data marked with the same letter are not significantly different by the LSD test at 5% level of significance.

An initial attempt to directly detect EaF82 in the pull-downed AKIN10 and EaF82 complex from extracts of TC transgenic floral tissues was unsuccessful. This could be due to low levels of such a complex present in the TC extracts with only limited EaF82 interacting with AKIN10. Since the *EaF82-S* without signal peptide was used for alkalinizing activity assay and in Y2H analysis, the EaF82-S peptide was spiked into both the vector control (C) and TC transgenic line extracts to perform co-IP performed assay. The idea was that if AKIN10 is indeed an interacting partner of EaF82, the added EaF82-S peptide should form a complex with endogenous AKIN10 in both lines, and by co-IP with anti-AKIN10 antibody, the complex should be able to pull-down. Consistent with this idea, we were able to pull-down AKIN10-EaF82-S complex from both the vector control and TC transgenic line extracts (Fig. 6b; lanes 4 and 7). Under the same incubation conditions without the anti-AKIN10 antibody, neither AKIN10 nor EaF82 was detected (Fig. 6b; lanes 3 and 6) while EaF82 was not detected when EaF82-S was incubating with protein A/G magnetic bead suspension (Fig. 6b; lane 8), indicating that no adsorption of AKIN/10 or EaF82-S to protein A/G magnetic beads occurred. The detected EaF82 (Fig. 6b; lane 4 and 7) could only be pull-downed by the anti-AKIN10 antibody bound protein A/G magnetic beads when it was present in the complex with AKIN10. These results confirm that AKIN10 can interact with EaF82, supporting the Y2H finding.

### Elevated levels of AKIN10 in *EaF82* transgenic flowers with ATP deficiency

In mammals, the level of AMPK is increased when the AMP/ATP ratio is high, resulting in phosphorylation of multiple downstream targets to increase ATP production and decrease ATP consumption^59^. In plants, the level of AKIN10 (AMPK homolog) was found to be also increased as the AMP/ATP ratio increased, while its transcriptional levels remained constant^60^. In addition, it has been reported that overexpression of *AKIN10* resulted in late flowering and defective silique development^61, 62^. For the identified EaF82 interacting partner AKIN10 (Fig. 6a, b), its gene *AKIN10* and another member *AKIN11* were detected in our RNAseq analysis but not differentially expressed (Supplementary Data Set 1). AKIN10 has also been reported to be degraded in a proteasome-dependent manner^63^ and induced by low energy conditions such as dark growth and hypoxia^60, 64^. To determine whether reduced ATP level and proteasome activity cause elevation of AKIN10 levels, we used immunoblotting with anti-AKIN10 antibody and discovered that TC-1 and TC-2 had an average of ∼4-fold higher levels of AKIN10 and ∼2-fold higher levels of AKIN11 compared to WT and vector control (Fig. 6c). These results are in agreement with the status of low ATP and reduced proteasome activity in transgenic lines (Fig. 5l, m).

## DISCUSSION

*EaF82* is a novel gene isolated from variegated ‘Golden Pothos’ plants with no known function^16^. It contains no intron and encodes a 120 amino acid long peptide^16^ with a 30 amino acid signal peptide at the *N*-terminus and four cysteine residues (Fig. 1a), sharing the typical features with numerous plant RALFs^20^. In present study, EaF82 identity as a member of RALF family was verified by its rapid alkalinization capacity (Fig. 1b, c), a hallmark of most RALFs^7^, and its overexpression affecting cellular pH regulation and cell wall modification related genes (Table 1; Supplementary Table 4). Phylogenetic analysis indicated that it belongs to clade IV (Fig. 1d), the least characterized group of RALFs. Clade IV members are mainly expressed in reproductive tissues^5^, but under certain conditions they may be induced in other tissues. For example, AtRALF8 reported to be abundant in pollen^13^ while it was also found to be induced in roots during drought and nematode infection^65^. *EaF82* expression was previously only found to correlate with IAA distribution in vegetative parts of ‘Golden Pothos’, a shy-flowering plant lacking flowers^15, 16^, but unknown in floral organs/tissues. Here, we demonstrated that ectopic expression of EaF82 in Arabidopsis delayed tapetum degeneration resulting in low pollen production and seed-setting, while appeared not to inhibit growth in vegetative parts (Fig. 5c), including seedlings (Supplementary Fig. 5a).

In flowering plants, pollen development occurs in anther with a complex process from initial microsporogenesis to the production of mature pollen grains^27, 66^. During this development, the sporophytic anther tissues, in particular the tapetum cell layer, play an essential role with the involvement of many genes to regulate developmental process, and provide materials for pollen wall formation^12, 66^. Recently, two tapetum-derived small peptides CASPARIAN STRIP INTEGRITY FACTOR 3 (CIF3) and 4 (CIF4) were shown to play an essential role in normal pollen wall deposition and tapetum function^67^. Our results also suggest that EaF82 may play an essential role in tapetum degeneration and pollen development. The histological analysis showed that the tapetum degeneration was delayed by EaF82 in TB lines (Fig. 3d; Supplementary Fig. 7). The involvement of EaF82 in tapetum degeneration and pollen development was further supported by our RNAseq and RT-qPCR results wherein we found that a group of *RALFs* reported in Arabidopsis tapetum mutants^12^ were also down-regulated in our transgenic lines. In addition, *CEP1* and seven additional protease genes identified in our down-regulated DEGs might be involved in impairing tapetum degeneration and causing reduction in nutrient supply to timely support pollen wall formation. Moreover, the down-regulation of seven *AGPs* important for pollen wall formation^37^ and many DEGs involved in nutrient transports (Supplementary Table 4) supports the observed pollen formation defect (Fig. 3d; Supplementary Fig. 7). In addition, many genes encoding PME and PMEI for cell wall modification^31^ and a group of auxin-responsive genes essential for pollen development^68^ were in down-regulated DEGs (Supplementary Table 4). Twenty-one transcription factors, including some known to regulate pollen formation and be pollen-specific, were also in down-regulated DEGs (Supplementary Table 4). All of these down-regulated genes lead us to believe that RALF-like peptide EaF82 compromised pollen formation via the impairment of tapetum degeneration.

Furthermore, our RNAseq and RT-qPCR data showed that the overexpression of *EaF82* resulted in the repression of seven *AtRALF* genes (*AtRALF4/8/9/15/19/25* and *At4g14020*) (Table 2). This intriguing phenomenon could be due to a feedback mechanism for regulating the expression of endogenous *AtRALFs*. Once these seven *AtRALF* propeptides undergo proteolytic processing to become mature peptides, they all have calculated molecular weights between 6.6 to 8.7 kD and pIs (isoelectric points) greater than 7 (9.3 - 10.6), indicating that they are basic small peptides (Table 2). Mature EaF82 (EaF82-S) also has a calculated pI of 8.7. It is possible that the overexpressed EaF82 peptide with higher accumulation levels and positive charge might compete with these AtRALFs to bind to their receptors and affect regulatory pathways leading to their transcriptional repression. Nevertheless, the regulatory mechanism of *EaF82* on the down-regulation of these *AtRALFs* needs future study.

Concerning the action mode of EaF82, the Y2H assay identified AKIN10 as its interacting partner (Fig. 6a). AKIN10 is a catalytic α-subunit of SnRK1 complex that comprises of an *N*-terminal Ser/Thr kinase domain, an adjacently linked ubiquitin-associated (UBA) domain, and a large *C*-terminal regulatory domain involved in the interaction with the regulatory (β and γ) subunits and upstream phosphatases^64^. *AKIN10* is expressed in flowers, anthers, and pollen^69, 70^. Under normal conditions, AKIN10 is localized mainly in cytosol^71^ where EaF82 is also localized^16^, supporting our detected interaction between EaF82 and AKIN10 in yeast (Fig. 6a). Our co-IP assay validated the interaction between EaF82 and AKIN10 (Fig. 6b). Concerning how EaF82 becomes a mature peptide and interacts with AKIN10, we speculate that EaF82 produced in one cell may be released into the extracellular space, at which then interacts with a receptor localized on cell surface of another cell to cross the plasma membrane and possibly its signal peptide is cleaved during or after its translocation of membrane. After getting into the cell cytosols, very likely the mature peptide (as EaF82-S rather than its nascent form) can then interact with kinase AKIN10. Nevertheless, further studies are necessary to understand the mechanism and possible effects of this interaction. Based on obtained results, it is reasonable to assume in EaF82 overexpressed lines that mature EaF82 binds to AKIN10 and modulates SnRK1 kinase activity and phosphorylation of downstream targets leading to transcriptomic changes. Although we did not examine any phosphorylation changes of proteins downstream of SnRK1, we did observe ∼5% transcriptomic changes at the early developmental stages of transgenic flowers overexpressing *EaF82*. Our Y2H results showed that EaF82 interacted with AKIN10 through its *C*-terminal end, which also contains kinase associated domain 1 (KA1, position 492-533) (Supplementary Fig. 11). The KA1 is a conserved domain of AMPKs from yeast to humans involved in autoinhibition, tethering of acidic phospholipids and binding of peptide ligands^72^. Whether EaF82 binds to the KA1 domain to affect SnRK1 activity needs further investigation as the function of KA1 domain in AKIN10 is still unclear^64^.

Our previous study indicated a correlation between the differentially expressed *EaF82* and the formation of color defective tissues in variegated leaves of *E. aureum*. We proposed that the light exposure upon developing leaves could drive the auxin flux unevenly causing differential expression of *EaF82* whose promoter is responsive to auxin^16^. However, whether the identified function of EaF82 inducing ATP deficiency could play a role in chloroplast biogenesis and then contribute leaf variegation formation is still inconclusive. In present study, the expression of EaF82 in Arabidopsis and tobacco plants did not cause variegation in leaves, at least indicating there are some other factors than EaF82 involving the variegated formation. Those factors might be lacking in both Arabidopsis and tobacco. The future study is warranted to determine whether EaF82 interacts with AKIN10, or other factors to cause leaf variegation formation.

## Materials and Methods

### Construction of genetic cassettes

Three pBI121 vector-based new genetic cassettes *EaF82p::EaF82-sGFP*, *35Sp::EaF82-sGFP* and *35Sp::EaF82* (designated as TA, TB and TC) were created in this study (Supplementary Fig. 2). The genetic cassette *EaF82p::GUS* was created in previous study^16^. The vector pBI121 carrying the *CaMV35S* promoter (*35Sp*) driving bacterial *uidA* (*GUS*) (*35Sp::GUS*) was used as a control (designated as C). To create these genetic cassettes, the intronless gene, *EaF82*, previously isolated from *E. aureum* containing a 1.2 kb *EaF82* promoter and a 0.8 kb full length *EaF82* including *EaF82* terminator^16^ was used. The two previously constructed plasmid DNAs, CEJ826 and CEJ937 (CEJ # is a nomenclature for plasmid DNA in Xie’s laboratory), and the pBI121 vector were used to construct these genetic cassettes. The plasmid DNA CEJ826 contains the *EaF82* promoter (*EaF82p*) and full length *EaF82* including *EaF82* terminator in pCR®-blunt 3.5 kb vector (Invitrogen). The plasmid DNA CEJ937 contains *35Sp* and 0.357 kb *EaF82* coding region without TGA stop codon in-frame fused with *sGFP* in pUC19 (Invitrogen). To construct the TC genetic cassette, the *EaF82* fragment isolated from CEJ826 by EcoRI-blunted and EcoRV digestion was ligated to vector pBI121 pre-digested by SacI-blunted and SmaI. The resultant contained *35Sp* driving full length *EaF82* including *EaF82* terminator. To construct the TB genetic cassette, the *35Sp::EaF82-sGFP-NosT* fragment isolated from CEJ937 by HindIII and EcoRI digestion was ligated to pBI121 pre-digested by HindIII and EcoRI. The resultant plasmid DNA was named CEJ964. To construct the TA genetic cassette, the *EaF82* promoter fragment isolated from CEJ826 by XbaI and EcoRV digestion was ligated to pre-digested CEJ964 by HindIII-blunted and XbaI. The resultant construct TA had *EaF82-sGFP* driven by *EaF82p*, whereas TB had the same *EaF82-sGFP* but driven by *35Sp*. The *Agrobacterium* strain LBA4404 was used to harbor these constructs.

### Created transgenic plants

Transgenic plants were generated with *A. thaliana* Col-0, *N. benthamiana* tobacco using previously described transformation methods^16, 73^ except the kanamycin concentration for selection in *N. benthamiana* was 300 mg/L. The Arabidopsis *EaF82p::GUS* transgenic plants were created previously^16^. All experiments were performed using homozygous lines.

### Alkalinization assay

The alkalinization assay was performed with tobacco (*N. tabacum,* W38) suspension-cultured cells (Supplementary Fig. 1b) derived from leaf calli following the method described by Pearce et al.^7^. The EaF82-S peptide (Fig. 1a) was synthesized by Lifetein (Somerset, NJ, USA) with a purity of 94.55%. Its molecular weight was 10 kD as confirmed by an Electrospray Ionization (ESI) Mass Spectrometry. Alkylated EaF82-S peptide was prepared as previously reported^7^ and used as a negative control. Initial 70 ml of suspension cells (OD_600_=0.2) in liquid medium, pH 5.6 containing MS basal salts and vitamins (Research Product International), were grown in a 300 ml flask under dark at 25°C with shaking at 160 rpm till the cell density reached 5-8 x 10^4^ cluster cells/ml after 3-4 days of subculture. For the assay, each 10 ml cell suspension was aliquoted into a 6-well plate and acclimated with shaking for 2-3 h. The peptide was reconstituted in ddH_2_O to give a concentration of 100 µM as a stock solution. Peptide concentrations of 1, 5, 10, 50 and 100 nM were used for assay. The pH was measured by a Mettler Toledo^TM^ InLab 413 pH Electrodes (Mettler-Toledo).

### RT-PCR and RT-qPCR

The procedures for RNA isolation, RT-PCR and RT-qPCR were the same as described in Hung et al.^14^. Data from three sets of biological samples were averaged. The primers are listed in Supplementary Table 9. The QuantumRNA™ Universal 18S Internal Standard (Invitrogen) was used as an internal control.

### RNAseq and DEG analyses

The Illumina sequencing was performed by DNA-Link (San Diego, CA, USA). The sequencing reactions were run on the Illumina NextSeq 500 with single-end 76 bp read. The Consensus Assessment of Sequence and Variation (CASAVA) software version 1.8.2 (Illumina) was used to remove adaptor sequences, nucleotide library indexes and generate fastq files. The RNAseq reads were mapped to the *A. thaliana* reference genome TAIR10 by TopHat^74^ to produce aligned reads and FPKM^75^. The gene annotations were based on NCBI database. The DEG analysis was conducted by Cufflinks and Cuffdiff^74^. The PANTHER classification system was used for functional pathway analysis^28^.

### Protein isolation, SDS-PAGE and immunoblotting

For extracting total proteins from leaves and seedlings, Plant Total Protein Extraction Kit (Sigma-Aldrich) was used. For extracting pollen proteins, the method was adopted from Chang & Huang^76^. In brief, hundreds of opened flowers were harvested in an ice-cold tube. About 5x volume of cold extraction buffer (HEPES potassium solution) was added. After brief mixing, the mixture was centrifuged at 350 *g* for 1 min to collect pollen. The purity of collected pollen was examined under a microscope (Supplementary Fig. 1c) before disruption by a blue pestle on ice. After 1 h incubation on ice, extracted pollen proteins in the supernatant were collected following centrifugation at 18,000 *g* for 30 min at 4°C. For extracting proteins from unopened flower clusters, tissues ground in liquid nitrogen were mixed in 1:2.5 ratio with extraction buffer containing 50 mM HEPES pH 7.8, 2 mM EDTA, 1 mM DTT and 1x Halt^TM^ Protease Inhibitor Cocktail, EDTA-free (Thermo Scientific). After centrifugation at 18,000 *g* for 20 min at 4°C, the protein extracts were collected.

SDS-PAGE, immunoblotting and the subsequent detection of chemiluminescent signals along with staining of blots were performed as previously described^16^. To detect EaF82, custom-made anti-EaF82 antibody was used^16^. To detect GFP, the blots were probed with 1:200 diluted mouse monoclonal anti-GFP (sc-9996, Santa Cruz Biotechnology). After three washes with TBST, blots were incubated with 1:10,000 diluted HRP-conjugated anti-mouse IgGκ (sc-516102, Santa Cruz Biotechnology). For detecting AKIN10, the blots were probed with 1:500 diluted anti-AKIN10 (AS10919, Agrisera) in PBST containing 1% dry-milk, followed by the 1:20,000 diluted HRP-conjugated anti-rabbit IgG (H+L) (AS014, ABclonal). For quantification of band intensity, the same blot was probed with 1:15,000 diluted plant actin mouse mAb (AC009, ABclonal) as an internal control, followed by the 1:20,000 diluted HRP-conjugated anti-mouse IgG (Jackson ImmunoResearch). Three independent experiments were performed and scanned images of band intensities on X-ray films were quantified using Image J (https://imagej.nih.gov/ij/). For examining the proteasome subunits, the same procedure used for detecting AKIN10 was also used except for the incubation conditions for each primary antibody. Anti-RPN6 (AS15 2832A, Agrisera) was 1:1,000 diluted in TBST containing 3% dry-milk and incubated at 23°C for 1 h. Anti-RPN10 (PHY0102S, PhytoAB) was 1:2,000 diluted in TBST and incubated at 23°C for 1 h. Anti-Rpt5a/b (PHY1747A, PhytoAB) was 1:1,000 diluted in PBST and incubated at 4°C for 16 h.

To determine the size of EaF82, the Novex™ tricine gel system (Invitrogen) for revealing low molecular weight proteins was used. In brief, extracted proteins were first mixed with equal volume of tricine SDS sample buffer containing 50 mM DTT. Before loading onto 16% tricine gel, the sample mixtures were denatured at 85°C for 2 min. The Spectra™ Multicolor Low Range Protein Ladder, a mixture of six proteins ranging from 1.7 to 40 kD, was used as size standards. Proteins were separated in tricine SDS running buffer under constant 50 volts for 4 h and later increasing to 100 volts for 1 h. They were then transferred to 0.2 μm PVDF membrane in Novex Tris-Glycine transfer buffer with 20% methanol under constant 20 volts for 90 min. Immunoblotting was performed as described for detecting EaF82.

### Histological analysis and seed counting

GUS assay was performed as previously described^16^. To detect GFP, tissues were observed directly under a fluorescence stereo microscope Nikon SMZ1000 (Nikon) equipped with ET-Narrow Band EGFP to minimize autofluorescence ex 480 nm/20 and em 510 nm/20 (49020, Chroma Technology). The images were taken by Nikon Digital Sight DS-Fi1 and analyzed by software NIS-ELEMENTS BR 3.0 (Nikon). To observe pollen, pollen grains were released by gently crushing the anthers on slides. For observing pollen germination, the method described by Krishnakumar & Oppenheimer^77^ was used to prepare slides containing germinated pollen grains. The pollen images were observed under a Zeiss LSM 710 microscope with ZEN software (Zeiss). For pollen viability assay, the stamens were tapped on a drop of iodine/potassium iodide TS 1 solution (RICCA chemical company) placed on a glass slide. After 5 min in dark, the pollen grains were imaged under a Keyence BZ-X700 microscope (Keyence). To observe pollen development, the whole unopened flower cluster was fixed in FAA solution (4% [v/v] formaldehyde, 5% [v/v] acetic acid and 50% [v/v] ethanol) with gentle vacuum for 5 min then kept at 4°C for 24 h. The dehydration, paraffin embedding, and sectioning steps were the same as described by Hung et al.^15^. After immobilizing on slides, the deparaffinized specimens were stained for 2-3 min with fresh 0.05% (w/v) Toluidine blue O solution in 0.1 M citrate phosphate buffer, pH 6.8 and then rinsed in running water. After mounting on Fluoromount (Sigma-Aldrich), the images were taken by a Keyence BZ-X700. For counting seeds, the images of seeds were counted by VisionWork®LS software (UVP).

### ATP measurement

The method of ATP measurement was adopted from Napolitano & Shain^78^. In brief, ATP was extracted by mixing 1 mg of tissue powder with 40 µl of 50 mM HEPES buffer, pH 7.4 containing 33 mAU/ml Novagen® proteinase K (Millipore). The mixture was incubated at 50°C for 15 min then at 80°C for 5 min. After centrifugation at 20,000 *g* for 10 min at 4°C, the supernatant was collected and used for ATP assay using an adenosine 5-triphosphate (ATP) bioluminescent assay kit (Sigma-Aldrich). The generated bioluminescence signal was measured with a SpectraMax M5 plate reader (Molecular Devices).

### Proteasomal activity assay

Proteasomal activity was assayed as described by Vallentine et al.^79^ except that 10 mM ATP and 5% (v/v) glycerol were included in the extraction buffer but omitted in the reaction buffer. Each assay had equal amounts of extracted proteins (3 µg). The generated signal was measured using a microplate reader PHERAstar (BMG Labtech).

### Yeast two-hybrid

The Y2H screening and 1-by-1 direct interaction assays were conducted by Hybrigenics Services SAS (Paris, France). The coding sequence for EaF82 (amino acid residues 30-120) was PCR-amplified from previously constructed CEJ982 containing *EaF82*, and cloned in frame with the LexA DNA binding domain (DBD) into pB27 vector as a *C*-terminal fusion to LexA (N-LexA-EaF82-C). Hybrigenics’ reference for this construct was hgx4998v1_pB27. The entire insert sequence in the construct was confirmed by sequencing and then used as a bait to screen a random-primed *A. thaliana* meiotic buds cDNA library constructed into pP6 vector. Cloning vectors pB27 and pP6 were derived from the original pBTM116^80^ and pGADGH^81^ plasmids, respectively.

The N-*LexA-EaF82*-C bait was tested in yeast and found neither toxic nor autoactivating by itself. Therefore, it was used for the ULTImate Y2H™ screening. A total of 115 million clones (11-fold the complexity of the library) were screened using a mating approach with YHGX13 (Y187 ade2-101::loxP-kanMX-loxP, mata) and L40ΔGal4 (mata) yeast strains as previously described^82^. A total of 271 His^+^ colonies were selected on a medium lacking tryptophan, leucine and histidine. The prey fragments of positive clones were amplified by PCR and sequenced at their 5’ and 3’ junctions. The obtained sequences were subjected to search the corresponding interacting protein in the GenBank database (NCBI) using a fully automated procedure. A confidence PBS score, which relies on both local and global scores, was assigned to each interaction as described by Formstecher et al.^83^. Briefly, the local score was analyzed by considering the redundancy and independency of prey fragments, as well as the distribution of reading frames and stop codons in overlapping fragments. Then, the global score was analyzed by taking into consideration of the interactions that were found in all the screens performed at Hybrigenics using the same library. The global score indicates the probability of a nonspecific interaction. The PBS scores were divided into six categories (A to E) for practical use purpose. Category A represents the highest confidence while D stands for the lowest confidence. Moreover, the category E specifically flags interactions involving highly connected prey domains previously found several times in screens performed on libraries derived from the same organism, while F stands for a false-positive with several of these highly connected domains that have been confirmed as false-positives of the technique. The PBS scores have been reported to correlate with the biological significance of interactions well^84, 85^. Interacting proteins with PBS scores A, B, C and D all have been confirmed to have biological relevance independently^83^. Therefore, all interacting proteins with PBS scores A, B, C and D could be considered as good candidates.

Using PBS score system, seven candidates were selected from the obtained EaF82 interacting proteins with score A, B, C or D (marked as P in Supplementary Table 7) to further perform 1-by-1 Y2H assay for validation. This direct 1-by-1 interaction assay was performed. Seven fragments were extracted from the ULTImate Y2H™ screening and cloned in frame with the Gal4 activation domain (AD) into plasmid pP7. The AD construct was checked by sequencing the 5’ and 3’ ends of the inserts. Hybrigenics’ references for these seven preys are listed in Supplementary Table 7.

To perform 1-by-1 pairwise Y2H interaction assays, the bait and prey constructs were transformed into the yeast haploid cells L40ΔGal4 (mata) and YHGX13 (Y187 ade2-101::loxP-kanMX-loxP, matα), respectively. These assays were based on the HIS3 reporter gene (growth assay without histidine). As negative controls, the bait plasmid was tested in the presence of empty prey vector (pP7) and all prey plasmids were tested with the empty bait vector (pB27). The interaction between SMAD and SMURF was used as the positive control^86^. Interaction pairs were tested in duplicate as two independent clones (clone 1 and clone 2) for the growth assay. For each interaction, undiluted and 10^-1^, 10^-2^, 10^-3^ dilutions of the diploid yeast cells (culture normalized at 5×10^7^ cells/ml) expressing both bait and prey constructs were spotted on several selective media. The DO-2 selective medium lacking tryptophan and leucine was used as a growth control to verify the presence of both the bait and prey plasmids. The different dilutions were also spotted on a selective medium without tryptophan, leucine and histidine (DO-3). Four different concentrations (1, 5, 10 and 50 mM) of 3-AT, an inhibitor of the HIS3 gene product, were added to the DO-3 plates to increase stringency and reduce possible autoactivation by the bait and prey constructs. The “DomSight” (Hybrigenics Services, SAS) displaying the comparison of the bait fragment and the Selected Interacting Domain (SID) of the prey proteins with the functional and structural domains (databases of protein domains: PFAM, SMART, TMHMM, SignalP, Coil algorithms) on these proteins were used for data visualization.

### Co-immunoprecipitation (co-IP) analysis

To validate the interaction between EaF82 and AKIN10, co-IP analysis was performed. Each 333 mg ground floral tissues from Arabidopsis vector control or TC transgenic line was resuspended in 1 ml of extraction buffer (50 mM Tris-HCl, pH 7.5 containing 150 mM NaCl, 0.05% Triton-X100, 10% glycerol, and both protease and phosphatase inhibitor cocktail) and kept on ice for 30 min for soaking. The samples were then homogenized using a BeadBug 6 microtube homogenizer (Benchmark Scientific) and centrifuged at 15,000 *g* for 10 min. The clear extracts were transferred to new tubes and centrifuged again to remove particulates. The clear extracts were transferred to new tubes.

To prepare the input samples, each 100 μl clear extracts was mixed with 37 μl of 4X LDS sample buffer (NuPAGE™, Invitrogen, USA) and 15 μl 10X reducing agent. The mixture was then heated at 70°C for 10 min and stored at −20°C until analysis. The remaining extracts were used for co-IP. First, both control and TC extracts were split into two 250 μl fractions. They then were spiked with or without 20 μg of EaF82-S peptide and divided into two tubes. After that, one tube was added 10 μg of anti-AKIN10 antibody (Cedarlane, USA) while the other tube received none (as a negative control). Another control was prepared by adding EaF82-S peptide alone to protein A+G magnetic beads. All samples were gently shaken overnight at 4°C for binding. After that, 50 μl of protein A/G magnetic beads pre-equilibrated with extraction buffer were added and incubated further for 3 h. After binding, all samples were centrifuged at 800 *g* for 1 min to separate beads. The supernatant was removed and the beads were first washed thrice with 400 μl of extraction buffer, followed by two washes with 0.1M Tris-HCl buffer, pH 7.5. The bound complex was released from beads by adding 150 μl of preheated (70°C) 4X LDS sample buffer and centrifuged at 10,000 *g* for 5 min. The eluates were then transferred to new tubes, mixed with 15 μl of 10X reducing agent (NuPAGE™, Invitrogen), and heated at 70°C for 10 min. Immunoblotting was then performed to detect EaF82 and AKIN10 in the eluate. Briefly, 20 μl of input and 30 μl of co-IP samples were loaded onto a 12% NuPAGE™ Bis-Tris gel (Invitrogen) and separated at 200 V for 40 min. Following separation, the proteins were transferred onto a PVDF membrane overnight at 4°C. The membrane was then blocked with fat-free milk powder in PBST (1%) for 1 h at room temperature, and then was first incubated with rabbit anti-AKIN10 antibody (1:500 dilution) followed by incubation with Veriblot IP detection reagent (Abcam, 1:500 in blocking solution). Protein bands were detected with Supersignal^TM^ west pico chemiluminescent substrate (Thermofisher) and images were captured with the iBright™ CL1500 imaging system (Thermofisher). To detect EaF82, the above blot was first stripped with Restore™ stripping buffer (Thermofisher), washed thrice for 10 min each with 1X PBST, and then blocked as described above. The remainder of the procedure to detect EaF82 was the same as described earlier.

### Peptide sequence analysis and phylogenetic tree construction

The SignalP 5.0 (http://www.cbs.dtu.dk/services/SignalP/)18 was used to predict the signal peptide. The Swiss Institute of Bionformatics Expasy server (https://web.expasy.org/compute_pi/)87 was used to predict pI and molecular weight. The construction of two phylogenetic trees was carried out following the information and procedures provided in Campbell & Turner^5^ except skipping a manual optimization; and MEGA X^22^ was used to perform evolutionary analysis of Arabidopsis clade IV-C RALFs with EaF82.

### Statistical analysis

Statistical analysis was performed with one-way ANOVA and Fisher’s least significant difference (LSD) test. The GraphPad Prism 7 (GraphPad Software) was used for calculating Michaelis–Menten constant (Km) and Vmax as well as statistic best fit value of R square and standard deviation of estimation (Sy.x).

### Data availability

Arabidopsis Genome Initiative locus identifiers for each gene mentioned in this study are listed in Tables 1 and 2, Supplementary Tables 2, 3, and 4, and Supplementary Data Sets 2 and 3. The accession number of *EaF82* is FJ666044. Raw data obtained from RNAseq analysis have been deposited into NCBI’s Gene Expression Omnibus under the accession codes GSE171459 (https://www.ncbi.nlm.nih.gov/geo/).

## Supporting information

All supplemental figures and tables

## Acknowledgements

We thank Qingping He and Maotao He for their assistance on histological analysis, Eva Johannes for the help on confocal microscopy, and Mythili Saravanan for the help on statistical analysis. This study was supported by the National Science Foundation grant (HRD-1400946 to J.X. and C.E.O.), National Institute of General Medical Sciences grant (SC1GM111178 to J.X.) and a Startup Fund of Golden LEAF Foundation to BRITE.

## Authors’ contributions

C.Y.H., J.X. and J.C. conceived and designed the experiments. C.Y.H., K.N.W., M.L.U., J.C., D.B.B., M.T., Q.Q., and J.Z. performed the experiments. C.Y.H., F.S.K., C.E.O., J.C., K.O.B. and J.X. analyzed the data. C.Y.H., F.S.K., and J.X. wrote the article with contributions of all the authors.

## Additional information

### 1. Supplementary Table 1, 5-9

Table S1. The summary of read counts of RNAseq analysis in current study.

Table S5. A subset of FPKM related to tapetum genes.

Table S6. The EaF82 interacting proteins.

Table S7. The Hybrigenics’ references of seven selected clones for 1-by-1 assays.

Table S8. Summary of interaction matrix and results.

Table S9. List of primer sequences used in current study.

### 2. Supplementary Figures 1-11

Fig. S1 Alkalinization assays of inactive EaF82-S as well as a representative of collected pollen grains and tobacco suspension cells.

Fig. S2. Genetic cassettes used in this study.

Fig. S3. Tissue specific expression of *EaF82* promoter in Arabidopsis transgenic *EaF82p::EaF82-sGFP* (TA) lines.

Fig. S4. Arabidopsis transgenic *EaF82p::EaF82-sGFP* lines (TA-1, -3, -4, and -5).

Fig. S5. Arabidopsis transgenic *35Sp::EaF82-sGFP* (TB) lines.

Fig. S6. RT-qPCR of *GUS* expression levels.

Fig. S7. Male gametophyte development of Arabidopsis transgenic *35Sp::EaF82-sGFP* (TB) line.

Fig. S8. The numbers of seeds per silique with different length (cm) in Arabidopsis transgenic 35Sp::EaF82 (TC) line.

Fig. S9. RNAseq analysis of early developmental flower buds of two independent *35Sp::EaF82* (TC1 and TC2) and vector control (C) transgenic lines.

Fig. S10. Solid growth tests on +/- Histidine and +/- 3-AT plates.

Fig. S11. A “DomSight” of AKIN10 (AT3G01090) displays the information of bait and prey structural, functional and interaction domains.

### Supplementary Tables 2-4 in Excel spreadsheets

Table S2. Common DEGs that both TC-1 and TC-2 increase two-fold compared to the vector control.

Table S3. Common DEGs that both TC-1 and TC-2 decrease two-fold compared to the vector control.

Table S4. A subset of upregulated and downregulated DEGs with FDR<0.05.

### Supplementary Data Sets 1-3 in Excel spreadsheets

Data Set S1. The FPKM values for normalized read counts of RNAseq analysis of Arabidopsis transgenic *35Sp::EaF82* (TC) and vector control (C) lines.

Data Set S2. Gene Ontology enrichment analysis of upregulated DEGs. Data Set S3. Gene Ontology enrichment analysis of downregulated DEGs.

