## Supplementary material for "A rapid alkalinization factor-like peptide EaF82 impairs tapetum degeneration during pollen development through induced ATP deficiency": All supplemental figures and tables: Combined Supplementary Tables - Figures 3-31-2023.pdf

**This Supplementary Data file includes:**

**1. Supplementary Table 1, 5-9**

Table S1. The summary of read counts of RNAseq analysis in current study.

Table S5. A subset of FPKM related to tapetum genes.

Table S6. The EaF82 interacting proteins.

Table S7. The Hybrigenics' references of seven selected clones for 1-by-1 assays.

Table S8. Summary of interaction matrix and results.

Table S9. List of primer sequences used in current study.

**2. Supplementary Figures 1-11**

Fig. S1 Alkalinization assays of inactive EaF82-S as well as a representative of collected pollen grains and tobacco suspension cells.

Fig. S2. Genetic cassettes used in this study.

Fig. S3. Tissue specific expression of *EaF82* promoter in Arabidopsis transgenic *EaF82p::EaF82-sGFP* (TA) lines.

Fig. S4. Arabidopsis transgenic *EaF82p::EaF82-sGFP* lines (TA-1, -3, -4, and -5).

Fig. S5. Arabidopsis transgenic *35Sp::EaF82-sGFP* (TB) lines.

Fig. S6. RT-qPCR of *GUS* expression levels.

Fig. S7. Male gametophyte development of Arabidopsis transgenic *35Sp::EaF82-sGFP* (TB) line.

Fig. S8. The numbers of seeds per silique with different length (cm) in Arabidopsis transgenic 35Sp::EaF82 (TC) line.

Fig. S9. RNAseq analysis of early developmental flower buds of two independent *35Sp::EaF82* (TC1 and TC2) and vector control (C) transgenic lines.

Fig. S10. Solid growth tests on +/- Histidine and +/- 3-AT plates.

Fig. S11. A “DomSight” of AKIN10 (AT3G01090) displays the information of bait and prey structural, functional and interaction domains.

**Supplementary Tables 2-4 in Excel spreadsheets**

Table S2. Common DEGs that both TC-1 and TC-2 increase two-fold compared to the vector control.

Table S3. Common DEGs that both TC-1 and TC-2 decrease two-fold compared to the vector control.

Table S4. A subset of upregulated and downregulated DEGs with FDR<0.05.

**Supplementary Data Sets 1-3 in Excel spreadsheets**

Data Set S1. The FPKM values for normalized read counts of RNAseq analysis of Arabidopsis transgenic *35Sp::EaF82* (TC) and vector control (C) lines.

Data Set S2. Gene Ontology enrichment analysis of upregulated DEGs.

Data Set S3. Gene Ontology enrichment analysis of downregulated DEGs.

**Table S1. The summary of read counts of RNAseq analysis in current study**

| Samples ID | Read | Read length bp | Index | Number of Reads |
|---|---|---|---|---|
| **C1** | SE | 76 | CTGAAGCT+AGGCTATA | 19,858,927 |
| **C2** | SE | 76 | TAATGCGC+AGGCTATA | 18,609,048 |
| **C3** | SE | 76 | CGGCTATG+AGGCTATA | 18,331,258 |
| **TC1-1** | SE | 76 | CTGAAGCT+GCCTCTAT | 20,035,018 |
| **TC1-2** | SE | 76 | TAATGCGC+GCCTCTAT | 19,216,782 |
| **TC1-3** | SE | 76 | CGGCTATG+GCCTCTAT | 20,693,114 |
| **TC2-1** | SE | 76 | CTGAAGCT+AGGATAGG | 23,037,461 |
| **TC2-2** | SE | 76 | TAATGCGC+AGGATAGG | 21,102,277 |
| **TC2-3** | SE | 76 | CGGCTATG+AGGATAGG | 21,339,999 |

**Table S5. A subset of FPKM related to tapetum genes.**

| Gene id | Gene short name | locus | length | FPKM_C1 | FPKM_C2 | FPKM_C3 | FPKM_TC1-1 | FPKM_TC1-2 | FPKM_TC1-3 | FPKM_TC2-1 | FPKM_TC2-2 | FPKM_TC2-3 |
|---|---|---|---|---|---|---|---|---|---|---|---|---|
| **Tapetum-related** | | | | | | | | | | | | |
| **AT5G07280** | **EMS1**/EXS | 5:2284829-2288855 | 4.03E+03 | 12.94 | 13.09 | 15.33 | 16.05 | 12.87 | 13.77 | 15.54 | 14.20 | 14.92 |
| **AT4G24972** | **TPD1** | 4:12837352-12840230 | 1.34E+03 | 6.83 | 4.42 | 6.49 | 3.74 | 5.79 | 5.30 | 5.00 | 4.87 | 4.98 |
| **AT1G34210** | **SERK2** | 1:12458563-12462905 | 2554 | 9.38 | 10.01 | 9.56 | 10.24 | 10.99 | 7.75 | 10.13 | 6.65 | 9.18 |
| **AT1G34210** | **SERK2** | 1:12458589-12462752 | 2267 | 6.15 | 5.44 | 8.14 | 5.18 | 5.28 | 4.95 | 5.58 | 10.70 | 6.98 |
| **AT1G71830** | **SERK1** | 1:27018157-27022117 | 2570 | 12.46 | 13.51 | 14.42 | 14.26 | 14.75 | 14.41 | 15.81 | 14.42 | 15.47 |
| **AT3G28470** | **TDF1**/MYB35 | 3:10674507-10675724 | 1025 | 1.37 | 2.02 | 1.91 | 2.62 | 0.97 | 1.34 | 2.08 | 1.48 | 2.71 |
| **AT2G16910** | **AMS** | 2:7331720-7334254 | 1893 | 33.39 | 33.40 | 45.57 | 48.79 | 29.50 | 33.23 | 47.66 | 35.97 | 44.85 |
| **AT5G56110** | **MYB188/MYB103/MYB80** | 5:22719190-22720664 | 963 | 3.39 | 2.56 | 4.60 | 2.55 | 1.20 | 2.07 | 2.87 | 1.67 | 3.67 |
| **AT5G22260** | **MS1** | 5:7367634-7370295 | 2.19E+03 | 2.59 | 2.50 | 3.25 | 3.28 | 2.23 | 1.66 | 3.51 | 2.43 | 3.18 |
| **AT4G21330** | **DYT1** | 4:11349921-11350694 | 624 | 0.13 | 0.68 | 0.28 | 1.92 | 0.27 | 0.87 | 1.58 | 1.09 | 1.32 |
| **AT4G20900** | **MS5** | 4:11184102-11185844 | 1353 | 0.00 | 0.00 | 0.00 | 0.00 | 0.00 | 0.00 | 0.00 | 0.00 | 0.00 |
| **AT4G20900** | **MS5** | 4:11184102-11185844 | 1305 | 1.17 | 0.73 | 1.80 | 1.81 | 0.57 | 1.00 | 1.29 | 1.38 | 1.47 |
| **AT3G51280** | **MS5-like** | 3:19037139-19039048 | 1649 | 20.20 | 27.64 | 26.55 | 38.31 | 41.52 | 32.95 | 50.05 | 41.90 | 41.58 |
| **AT3G11980** | **MS2**/FAR2 | 3:3814230-3817117 | 2117 | 51.40 | 33.34 | 54.30 | 60.38 | 45.99 | 35.70 | 64.57 | 52.92 | 69.76 |

| | | | | | | | | | | | | |
|---|---|---|---|---|---|---|---|---|---|---|---|---|
| **AT4G33790** | **MS2-like/**FAR3 | 4:16204095-16207956 | 1956 | 0.73 | 0.49 | 1.02 | 1.31 | 0.54 | 1.30 | 0.22 | 1.51 | 0.75 |
| **AT4G33790** | **MS2-like/**FAR3 | 4:16204095-16207956 | 1776 | 39.84 | 34.90 | 39.31 | 37.02 | 40.67 | 27.61 | 28.08 | 28.10 | 32.53 |
| **AT4G27330** | **SPL/NZZ** | 4:13682077-13683594 | 1329 | 6.96 | 6.80 | 9.52 | 5.98 | 3.94 | 6.38 | 5.31 | 5.33 | 5.67 |
| **AT5G50260** | **CEP1*** | 5:20455316-20457068 | 1580 | **6.55** | **2.78** | **5.29** | **0.27** | **0.37** | **0.53** | **0.28** | **0.15** | **0.22** |
| **AT2G01570** | **RGA** | 2:255246-257568 | 2.32E+03 | 33.43 | 39.86 | 40.28 | 38.23 | 40.33 | 38.79 | 37.86 | 40.58 | 43.17 |
| **AT4G02780** | **GA1** | 4:1237670-1244822 | 2675 | 0.41 | 0.62 | 0.57 | 0.59 | 0.64 | 0.38 | 0.56 | 0.62 | 0.70 |
| **AT1G14920** | **GAI** | 1:5148981-5151415 | 2434 | 19.30 | 23.02 | 21.23 | 25.44 | 28.62 | 24.79 | 25.76 | 24.24 | 29.28 |
| **AT5G06100** | **MYB33** | 5:1837834-1840706 | 2266 | 2.96 | 2.58 | 4.12 | 3.55 | 3.76 | 6.08 | 6.83 | 5.50 | 2.60 |
| **AT5G06100** | **MYB33** | 5:1837834-1840727 | 2206 | 8.33 | 7.79 | 6.27 | 8.20 | 6.43 | 4.91 | 4.10 | 5.46 | 8.46 |
| **AT5G06100** | **MYB33** | 5:1837895-1840701 | 2537 | 0.00 | 0.00 | 0.02 | 0.00 | 0.00 | 0.00 | 0.00 | 0.00 | 0.13 |
| **AT5G06100** | **MYB33** | 5:1837913-1840159 | 1741 | 0.00 | 0.00 | 0.00 | 0.00 | 0.21 | 0.00 | 0.23 | 0.39 | 0.73 |
| **AT5G06100** | **MYB33** | 5:1837913-1840706 | 2443 | 0.20 | 0.00 | 1.79 | 0.62 | 0.72 | 1.02 | 0.36 | 0.40 | 0.04 |
| **AT5G06110** | **MYB33** | 5:1840754-1843872 | 2554 | 28.83 | 19.05 | 32.29 | 36.99 | 33.59 | 28.20 | 35.01 | 26.77 | 31.72 |
| **AT3G11440** | **MYB65** | 3:3602035-3605172 | 2.23E+03 | 0.00 | 0.25 | 0.26 | 0.00 | 0.00 | 0.00 | 0.00 | 0.00 | 0.00 |
| **AT3G11440** | **MYB65** | 3:3602035-3605172 | 2140 | 0.00 | 1.94 | 0.63 | 0.42 | 1.31 | 0.00 | 0.37 | 0.00 | 0.78 |
| **AT3G11440** | **MYB65** | 3:3602348-3605110 | 2089 | 0.00 | 0.00 | 0.00 | 0.00 | 0.00 | 0.00 | 0.00 | 0.00 | 0.00 |
| **AT3G11440** | **MYB65** | 3:3602883-3605104 | 2009 | 3.24 | 0.53 | 2.97 | 2.36 | 1.68 | 3.34 | 3.44 | 3.07 | 2.28 |

*: The gene is in the downregulated DEG list.

### Table S6. The EaF82 interacting proteins

| | name | ATG | # clones | score |
|---|---|---|---|---|
| | Arabidopsis thaliana - BIP1 | AT5G28540 | 16 | A |
| **p** | Arabidopsis thaliana - ALATS | AT1G50200 | 10 | A |
| **p** | Arabidopsis thaliana - SYTA | AT2G20990 | 10 | A |
| | Arabidopsis thaliana - F17H15.1 | AT2G25970 | 7 | A |
| | Arabidopsis thaliana - MNJ7.18 | AT5G47590 | 7 | A |
| | Arabidopsis thaliana - SOX | AT3G01910 | 7 | B |
| | Arabidopsis thaliana - F14O23.22 | AT1G71840 | 5 | B |
| | Arabidopsis thaliana - HSC70-1 | AT5G02500 | 5 | B |
| | Arabidopsis thaliana - F5F19.6 | AT1G52000 | 4 | B |
| | Arabidopsis thaliana - FP3 | AT5G63530 | 4 | B |
| **p** | Arabidopsis thaliana - PAPP2C | AT1G22280 | 4 | B |
| **p** | Arabidopsis thaliana - ABCF4 | AT3G54540 | 3 | C |
| | Arabidopsis thaliana - LOS2 | AT2G36530 | 3 | C |
| | Arabidopsis thaliana - SPDS2 | AT1G70310 | 3 | C |
| | Arabidopsis thaliana - emb1507 | AT1G20960 | 3 | C |
| | Arabidopsis thaliana - heat shock protein 70-3 | AT3G09440 | 3 | C |

| | | | | |
|---|---|---|---|---|
| | Arabidopsis thaliana - MFB16.14 | AT5G50740 | 5 | D |
| | Arabidopsis thaliana - PATL1 | AT1G72150 | 5 | D |
| | Arabidopsis thaliana - ACT2 | AT3G18780 | 2 | D |
| | Arabidopsis thaliana - AMY3 | AT1G69830 | 2 | D |
| | Arabidopsis thaliana - Carbohydratebinding-like fold | AT3G62360 | 2 | D |
| | Arabidopsis thaliana - F13M22.2 | AT2G37520 | 2 | D |
| **p** | Arabidopsis thaliana - FKBP-like | AT3G55520 | 2 | D |
| | Arabidopsis thaliana Translation elongation factor EFG/EF2 protein (SCO1) | AT1G62750 | 2 | D |
| | Arabidopsis thaliana - HIP1 | AT4G22670 | 2 | D |
| | Arabidopsis thaliana - IDM2 | AT1G54840 | 2 | D |
| | Arabidopsis thaliana - MYC3 | AT5G46760 | 2 | D |
| | Arabidopsis thaliana - SK5 | AT3G60020 | 2 | D |
| | Arabidopsis thaliana - SRC2 | AT1G09070 | 2 | D |
| **p** | Arabidopsis thaliana - TCH4 | AT5G57560 | 2 | D |
| | Arabidopsis thaliana - clathrin | AT5G05010 | 2 | D |
| **p** | Arabidopsis thaliana - AKIN10 | AT3G01090 | 1 | D |
| | Arabidopsis thaliana - DRH1 | AT3G01540 | 1 | D |
| | Arabidopsis thaliana - F18O14.16 | AT1G19400 | 1 | D |
| | Arabidopsis thaliana - HSP91 | AT1G79930 | 1 | D |
| | Arabidopsis thaliana - MBP1 | AT1G52040 | 1 | D |
| | Arabidopsis thaliana - MDJ22.18 | AT5G22760 | 1 | D |
| | Arabidopsis thaliana - MJJ3.25 | AT5G05830 | 1 | D |
| | Arabidopsis thaliana - MPK17.2 | AT5G36740 | 1 | D |
| | Arabidopsis thaliana - PBB2 | AT5G40580 | 1 | D |
| | Arabidopsis thaliana - Phox3 | AT5G20360 | 1 | D |
| | Arabidopsis thaliana - SGT1A | AT4G23570 | 1 | D |
| | Arabidopsis thaliana - T19D16.19 | AT1G10890 | 1 | D |
| | Arabidopsis thaliana - T1K7.11 | AT1G26520 | 1 | D |
| | Arabidopsis thaliana - cpHsc70-1 | AT4G24280 | 1 | D |
| | Arabidopsis thaliana - hypothetical protein | AT3G59430 | 1 | D |

**Table S7. The Hybrigenics' references of seven selected clones for 1-by-1 assays**

| |
|---|
| ATMB_RP1_hgx4998v1_pB27_A-237 ABCF4 [402-557][*] |
| ATMB_RP1_hgx4998v1_pB27_A-240 ALATS [36-282] |
| ATMB_RP1_hgx4998v1_pB27_A-66 FKBP-like peptidyl-prolyl cis-trans isomerase family protein [43-190] |
| ATMB_RP1_hgx4998v1_pB27_A-262 PAPP2C [177-287] |
| ATMB_RP1_hgx4998v1_pB27_A-94 TCH4 [24-178] |
| ATMB_RP1_hgx4998v1_pB27_A-125 AKIN10 [422-535] |
| ATMB_RP1_hgx4998v1_pB27_A-157 SYTA [309-536] |

[*]The amino acid residues of proteins been identified in prey fragments are indicated in brackets. The prey fragments that do not stop with the natural stop codon of the protein have a short additional peptide at the C-terminal end which comes from the SfiI restriction side, being used for cloning of the library fragments.

### Table S8. Summary of interaction matrix and results

| Interaction Matrix | | | | | Selection Medium | | | | | |
|---|---|---|---|---|---|---|---|---|---|---|
| N# Interaction* | Type | Bait | Prey | Prey clone reference | DO-2 | DO-3 | DO-3 + 1mM 3-AT | DO-3 + 5mM 3-AT | DO-3 + 10mM 3-AT | DO-3 + 50mM 3-AT |
| 1 | Hybrigenics' positive control | SMAD | SMURF | / | + | + | + | + | + | + |
| 2 | Negative control | F82 | pP7ø | / | + | - | - | - | - | - |
| 3 | Negative control | pB27ø | ABCF4 | pB27_A-237 | + | - | - | - | - | - |
| 4 | Interaction | F82 | ABCF4 | pB27_A-237 | + | + | + | + | +/- | - |
| 5 | Negative control | pB27ø | ALATS | pB27_A-240 | + | - | - | - | - | - |
| 6 | Interaction | F82 | ALATS | pB27_A-240 | + | + | + | - | - | - |
| 7 | Negative control | pB27ø | FKBP_like peptidyl-prolyl cis-trans isomerase family protein | pB27_A-66 | + | - | - | - | - | - |
| 8 | Interaction | F82 | FKBP_like peptidyl-prolyl cis-trans isomerase family protein | pB27_A-66 | + | + | + | +/- | - | - |
| 9 | Negative control | pB27ø | PAPP2C | pB27_A-262 | + | - | - | - | - | - |
| 10 | Interaction | F82 | PAPP2C | pB27_A-262 | + | + | + | + | + | |
| 11 | Negative control | pB27ø | TCH4 | pB27_A-94 | + | - | - | - | - | - |
| 12 | Interaction | F82 | TCH4 | pB27_A-94 | + | + | + | - | - | - |
| 13 | Negative control | pB27ø | AKIN10 | pB27_A-125 | + | - | - | - | - | - |
| 14 | Interaction | F82 | AKIN10 | pB27_A-125 | + | + | + | + | +/- | - |
| 15 | Negative control | pB27ø | SYTA | pB27_A-157 | + | - | - | - | - | - |
| 16 | Interaction | F82 | SYTA | pB27_A-157 | + | + | + | - | - | - |

*A single clone of each negative control is spotted in duplicates. pB27: LexA DNA Binding Domain (DBD) vector (LexA-bait); pB27ø: empty pB27 vector; pP7/pP6: Gal4 Activation Domain (AD) vector (AD-prey); pP7ø: empty pP7 vector; DO-2: selective medium without tryptophan and leucine; DO-3: selective medium without tryptophan, leucine and histidine; 3-AT: 3-aminotriazole.

### Table S9. List of primer sequences used in current study

| AGI ID | gene name | Forward sequence (5'-3') | Reverse sequence (5'-3') |
|---|---|---|---|
| **At1G19890** | *MGH3* | AGGCAGCCGAAGCATACTTG | ACCCTTTTGGCGTGAATCG |
| **At1G19960** | *AT1G19960* | CGGCGGCTCCTTTGG | CCACACTTGCTTCCTCACCTTT |
| **At1G21000** | *AT1G21000* | ACGGTCCCAATGGTGATCA | GGAGAGAAACTAAAACCACCTTCATC |
| **At1G24520** | *BCP1* | CGATGACGATGCAGCTCCTA | TAGAGGACCAGCCACTGCAA |
| **At1G28270** | *RALFL4* | CAATGCAACCTACCCTTTAACCA | GTCTTCCCCGATGCAACCT |
| **At1G29140** | *AT1G29140* | TGAGTGCAAGGGTCGTGAGA | TTGTCGGTCACGGCTTCTTT |
| **At1G35490** | *AT1G35490* | CATCGAGCGCGTTTGAGA | TTGTAGCACTTGGATGGTCCTTT |
| **At1G61563** | *RALFL8* | CCCACCCTAACACCTGCAA | CTTTGGACCTGTTTCACGATGG |
| **At1G61566** | *RALFL9* | CCCACCCTAACACCTGCAA | ATCTTCTCGCATCCTCTCTGGTA |
| **At1G80660** | *HA9* | TGGCACCGTGTCAGCAAA | CTTGCATCCTCCCGAAGGT |
| **At2G07040** | *PRK2A* | GCGCGCAAGAGCTCATG | TTGGTCACACGGCTTTGTTTT |
| **At2G07560** | *HA6* | GGGTTGCACGTCAGAGAGTT | TTGGAGGGTCAAAGAGTGGC |

| | | | |
|---|---|---|---|
| **At2G18470** | *PERK4* | GCAAGAAATGGCTCGAATGG | CGCCCCGAATGACGAAT |
| **At2G21480** | *AT2G21480* | TGAACGGTGTCGAGGTCTTG | ATCCCCTGCTTCCCCATACT |
| **At2G22055** | *RALFL15* | CAAAGCCCACCCTAACACCT | ACCTGTATCACGACGGCAAC |
| **At2G33775** | *RALFL19* | TGTTGATCCTTGGCCTCTTGA | GGGTCCATGTGGCATTGG |
| **At2G36080** | *ABS2* | GGAGAGCCAAAGAGGCAACT | CCGAATCTAGCTGGCACTCC |
| **At2G45660** | *AGL20* | ACTCTTGGGAGAAGGCATAGGAA | TCAAGCTGTTGCTCAATCTGTTG |
| **At3G08560** | *VHA-E2* | AAGCCAAAGTCGGGTCTCCTA | AGGAGGTGGAGGGAGGAAGA |
| **At3G25165** | *VHA-E2* | GGACGAGTTGAGGCTAACGATAA | GGAGCATTAGGACGCAAGCA |
| **At3G42640** | *HA8* | GGGGAGCAAGAGGCTTCAAT | CCTTCGAGGAGACGAGCATC |
| **At4G10603** | *AT4G10603* | CCGTTGTTCGGCTAGGTGTAAG | CATTGCCTTGCCCCCTTTA |
| **At4G14020** | *AT4G14020* | ACATTTACCGAAAACGACGAAGAA | TTCTTTCACCCACGTGTTATTCA |
| **At4G24540** | *AGL24* | ACGTTGGAAAGGGCAAAACTG | TTTGTGGTCACCGACTCTGTCT |
| **At4G35900** | *FD* | GCAAGACTCAAGAGACAACAAG | CAAAATGGAGCTGTGGAAGAC |
| **At4G39110** | *AT4G39110* | AGGGCAAAGCAGAAGAGACC | TTCGTTGGTGGTGACTGAGG |
| **At5G17480** | *PC1* | ATGGCTGATGCAACGGAGAA | GAGCTTCTTCGAGTTCGGCT |
| **At5G28680** | *ANX2* | AGCAGTTTTTGTCTTGTTGGGG | CTGGAGGCTTGTTTCTGCAA |
| **At5G45880** | *AT5G45880* | CTTGCCGTGTCCAATTCGTT | TTCTGCTCCTGCACTCCAAC |
| **At5G57350** | *HA3* | TTCCTGCTGATGCACGTCTT | TTCCCCTGGACCTTTCGTTG |
| **At5G62165** | *AGL42* | CCAGCAATCACGACTCACAAAT | TTGTTATCATGTGGCTTGCTTCTT |
| | *EaF82* | GCATGTGGTGTCACTCCTCTTTAT | ATCTTCTTCCGCAGTCCCAAA |
| | *NPTII* | AAGATGGATTGCACGCAGGTTC | ACGGGTAGCCAACGCTATGTC |
| | *GUS2* | TGGCCTGGCAGGAGAAACT | ACGTATCCACGCCGTATTCG |

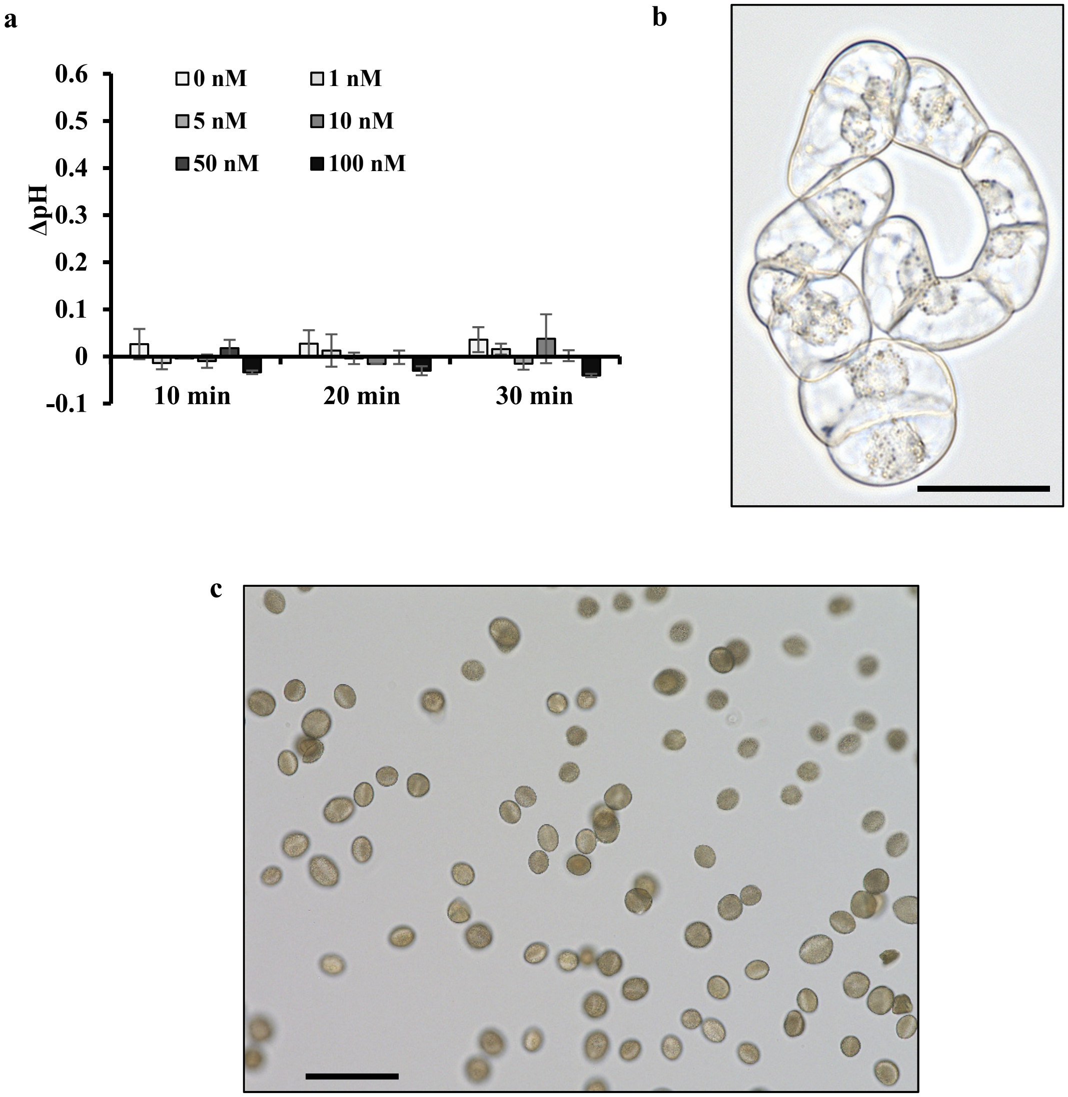


**Fig. S1** Alkalinization assays of inactive EaF82-S as well as a representative of collected pollen grains and tobacco suspension cells. **a** Alkalinization assays of inactive EaF82-S. Data plotted is the concentration curve. The pH changes (Δ pH) were calculated as measured pH value after applying inactive EaF82-S for 10, 20 and 30 min minus the pH value at 0 min at each concentration from 0 to 100nM. The data plotted were the average of three independent experiments ± SD. **b** A representative of tobacco suspension cells used for alkalinization assay. Bar=50μm. **c** A representative of collected pollen in HEPES potassium salt buffer for protein isolation. Bar=100μm.

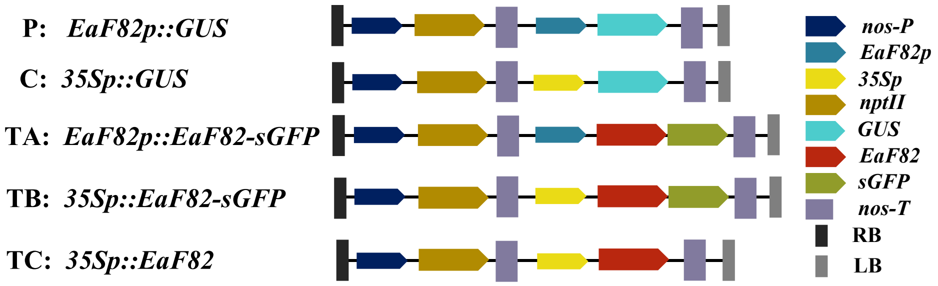


**Fig. S2** Genetic cassettes used in this study. *EaF82p: EaF82* promoter; *35Sp*: CaMV *35S* promoter; *sGFP: Green Fluorescent Protein* (S65T)*; GUS*: *uidA*; *nptII: neomycin phosphotransferase II*; *nos-P: nopaline synthase* promoter; *nos*-T: *nos* terminator; RB: right border; LB, left border.

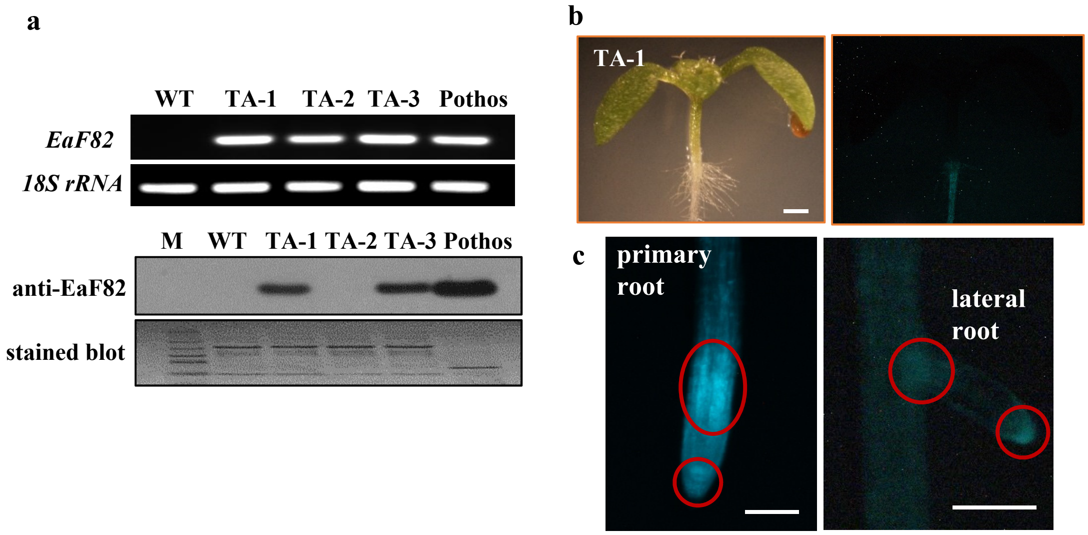


**Fig. S3** Tissue specific expression of *EaF82* promoter in Arabidopsis transgenic *EaF82p::EaF82-sGFP* (TA) lines. (**a**) RT-PCR (upper panel) and immunoblot (lower panel) of three 7-day old subline seedlings (TA-1, 2, and 3). PCR was performed using primer pairs specific to *EaF82* and *18S rRNA* gene (as internal control). Stained blot shows protein loading. *EaF82* originated from 'Golden Pothos' was used as a positive control. (**b**) 10-day old light grown seedlings (left) and the detected GFP signals in roots (right). Bar = 1mm. (**c**) GFP signals detected in primary and lateral roots are circulated in red. Bar = 500 µm.

a

b

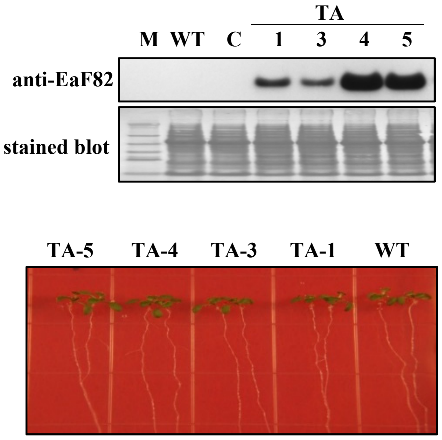


**Fig. S4** Arabidopsis transgenic *EaF82p::EaF82-sGFP* lines (TA-1, -3, -4, and -5). (**a**) Immunoblot of seedling proteins against anti-EaF82. C: vector control. WT: wild-type. M: protein size marker. (**b**) 10-day old seedlings germinated on MS medium show normal growth as WT.

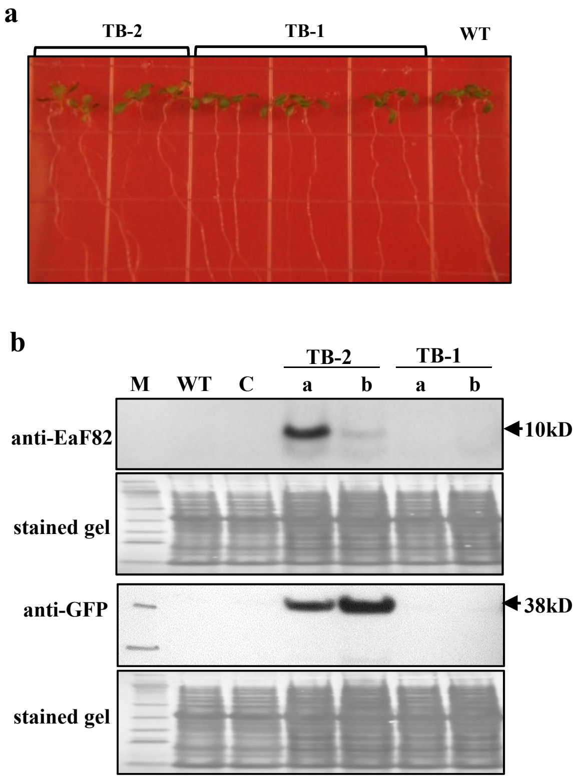


**Fig. S5** Arabidopsis transgenic *35Sp::EaF82-sGFP* (TB) lines. (**a**) 10-day old seedlings germinated on MS medium show normal growth as wild-type (WT). (**b**) Immunoblot of seedling proteins against anti-EaF82 and anti-GFP. TB-1 and -2 are two independent lines, while a, and b are two sublines.

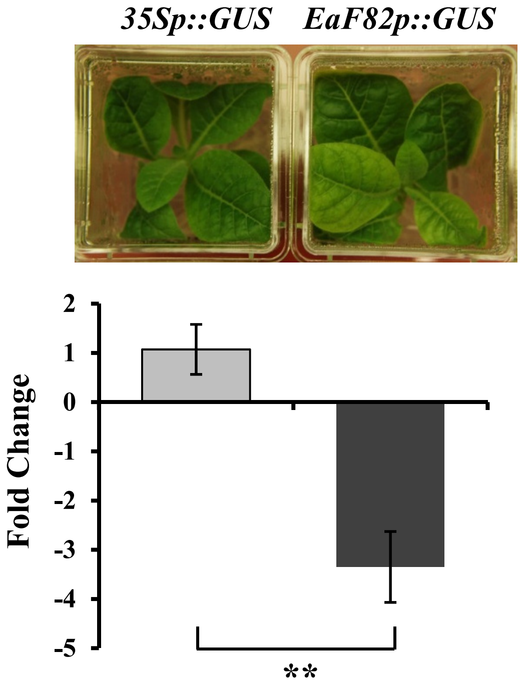


**Fig. S6** RT-qPCR of *GUS* expression levels. Transgenic tobacco plants carrying *GUS* driven by CaMV *35S* promoter (*35Sp:GUS*) and *EaF82* promoter (*EaF82p::GUS*), respectively (upper panel). GUS expression levels of each transgenic line were first normalized with their internal *18S rRNA* expression level. The fold changes were then calculated as *EaF82p::GUS* comparing to *35Sp::GUS* (as 1). Data plotted are the average of fold changes in three biological replicates ± SD. ** $P<0.01$.

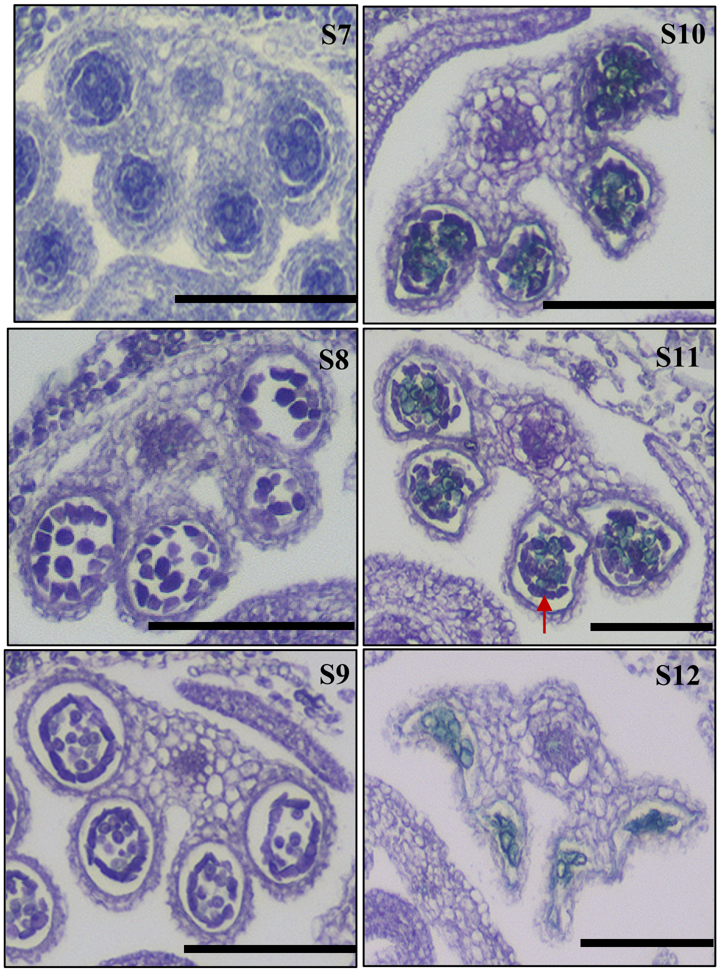


**Fig. S7** Male gametophyte development of Arabidopsis transgenic *35Sp::EaF82-sGFP* (TB) line. The stages S7 to S12 of a second independent TB flower cluster. The delayed degeneration of tapetum at stage S11 is indicated in red arrow. Underdeveloped pollen at stage S12 are stained in light green. Bar = 100 μm.

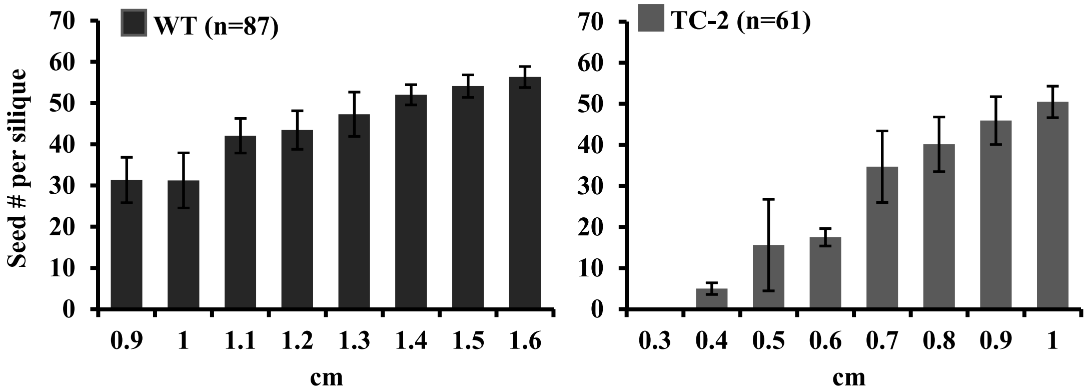


**Fig. S8** The numbers of seeds per silique with different length (cm) in Arabidopsis transgenic 35Sp::EaF82 (TC) line. Data plotted are the average ± SD. WT: wild-type; n: number of siliques.

**a**

| Genomes | Total listed | Non-detected | % of non-detected |
|---|---|---|---|
| **nuclear** | **32541** | **4382** | **13.5** |
| **mitochondrial** | **152** | **13** | **8.6** |
| **plastid** | **133** | **16** | **12.0** |

**b**

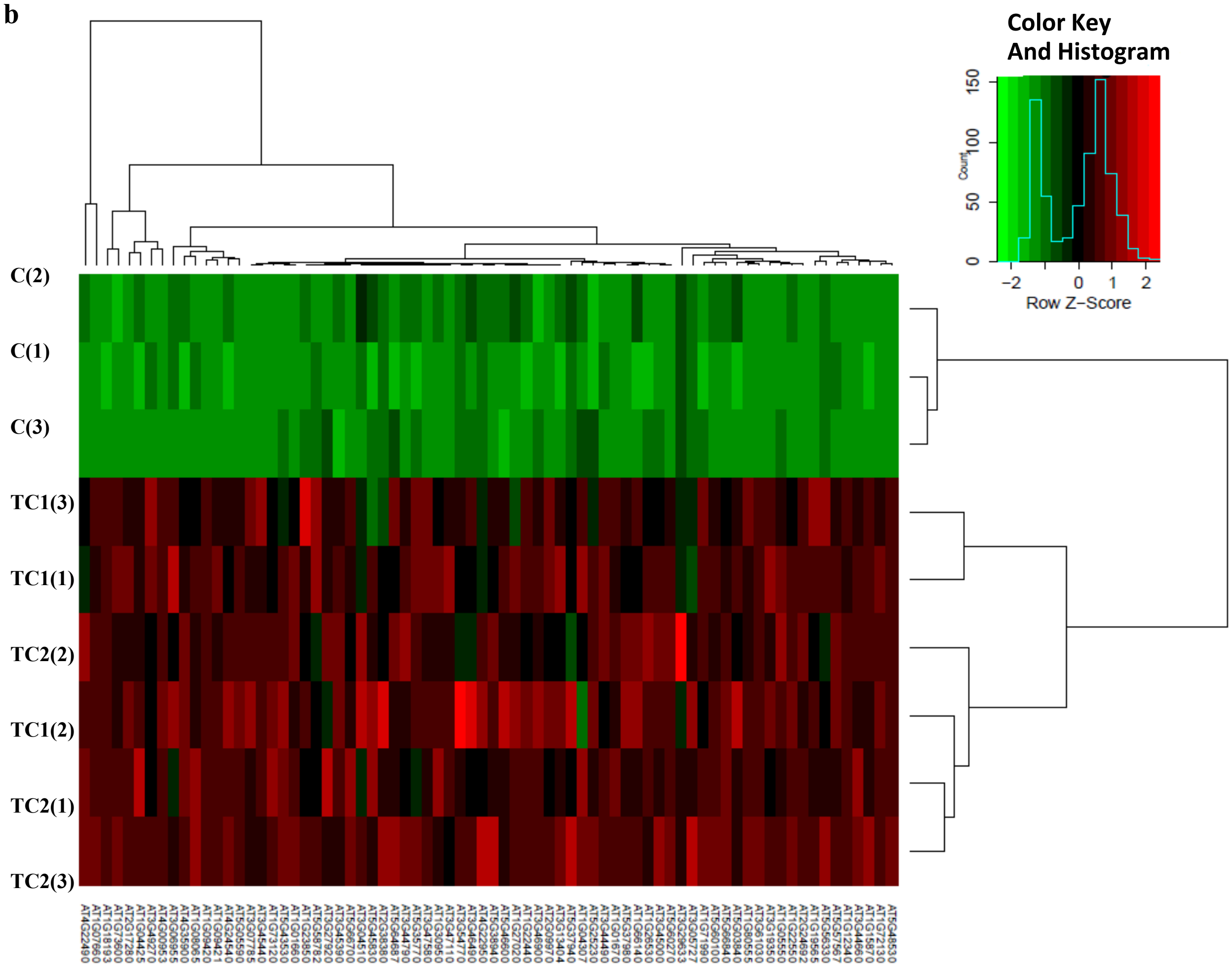


**Fig. S9** RNAseq analysis of early developmental flower buds of two independent *35Sp::EaF82* (TC1 and TC2) and vector control (C) transgenic lines. (**a**) The numbers of FPKM_gene listed in Data Set 1. (**b**) Heat map of hierarchical cluster analysis on DEGs of TC1 and TC2 both with $\log_2(FC) \leq -1.5$. Three biological replicates are indicated as TC1 (1, 2 and 3), TC2 (1, 2 and 3) and C (1, 2 and 3).

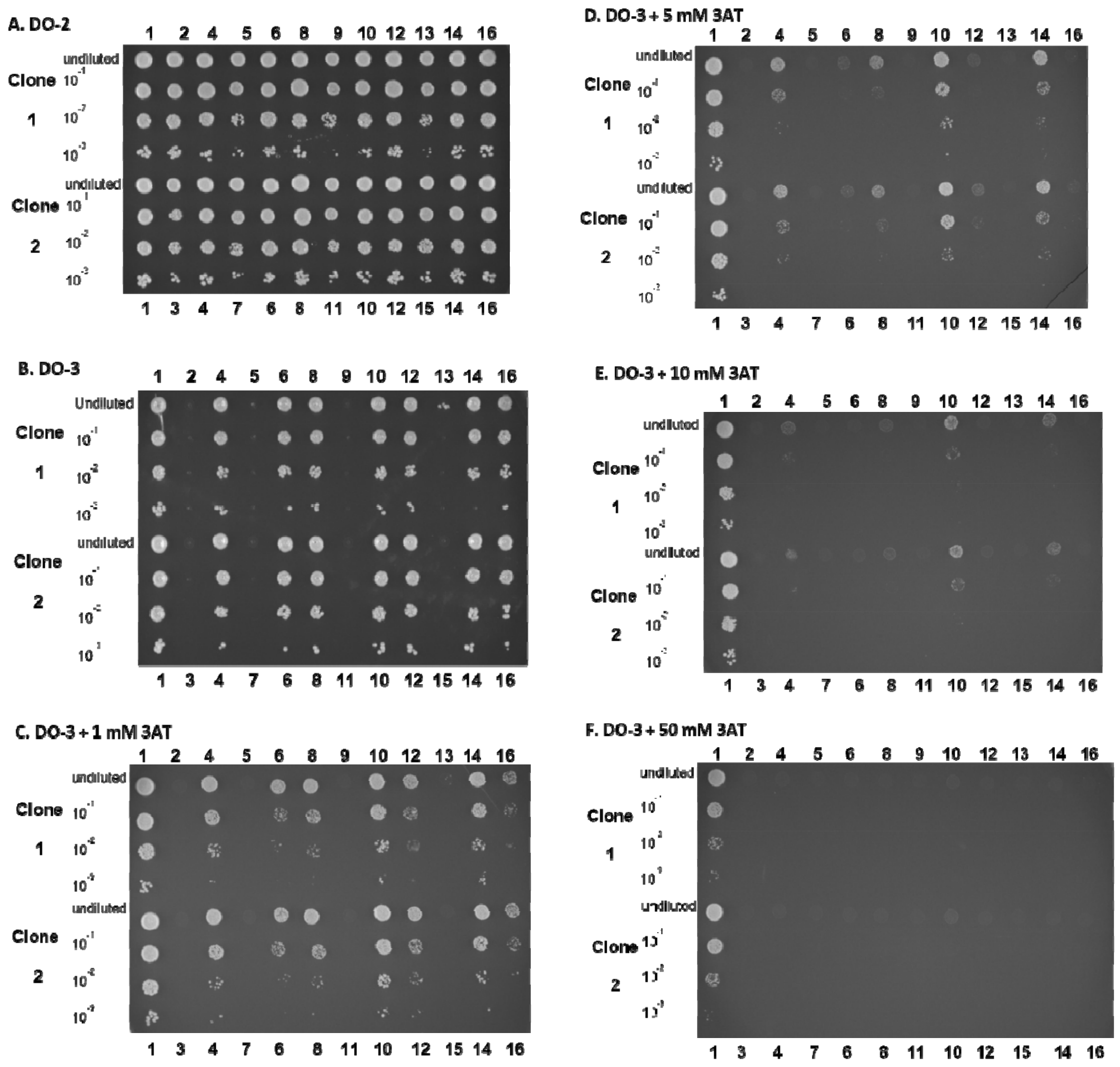

**Fig. S10** Solid growth tests on +/- Histidine and +/- 3-AT plates. **A** DO-2, **B** DO-3, **C** DO-3 + 1 mM AT, **D** DO-3 + 5 mM AT, **E** DO-3 + 10 mM AT, and **F** DO-3 + 50 mM AT. Detailed information of the interacting #1-16 are listed in Supplementary Table 8.

**DomSight index**

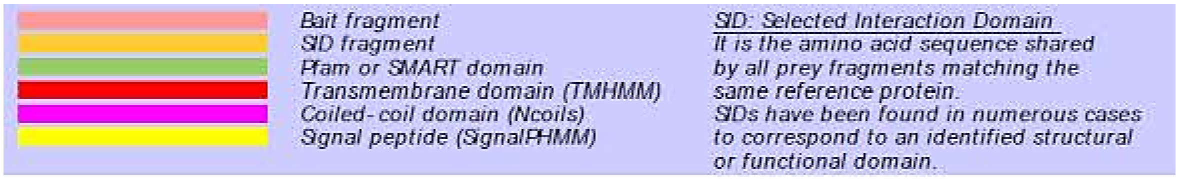


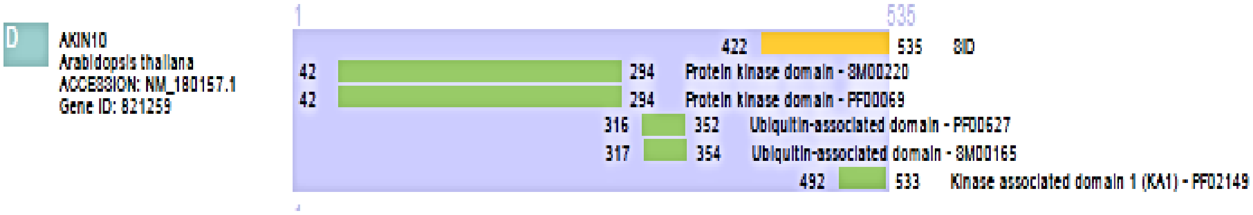


**Fig. S11** A "DomSight" of AKIN10 (AT3G01090) displays the information of bait and prey structural, functional and interaction domains. The SID domain (amino acid residues 422-535) contains kinase associated domain 1 (KA1).
